# Dissecting translation elongation dynamics through ultra-long tracking of single ribosomes

**DOI:** 10.1101/2024.04.08.588516

**Authors:** Maximilian F. Madern, Sora Yang, Olivier Witteveen, Marianne Bauer, Marvin E. Tanenbaum

**Author notes:** denotes equal contribution (Maximilian F. Madern, Sora Yang).

## Abstract

mRNA translation by ribosomes is a highly dynamic and heterogeneous process. However, current approaches cannot readily resolve individual ribosomes during translation, limiting our understanding of translation dynamics. Here, we develop an imaging approach based on Stopless-ORF circular RNAs (socRNAs) to monitor individual translating ribosomes for hours. Using the socRNA imaging technology we obtained accurate measurements of ribosome pausing on various problematic RNA sequences or induced by ribosome-targeting drugs. In addition, we identified a novel translation factor involved in translation elongation, and revealed that translocation rates of ribosomes vary, indicative of intracellular ribosomal heterogeneity. Finally, socRNAs allow very sensitive measurements of translation elongation fidelity, revealing widespread frameshifting during translation. In summary, our single-ribosome imaging approach provides a detailed view of ribosome translocation kinetics and a powerful new tool to study the translation elongation phase.

## Introduction

mRNA translation by the ribosome is a key step in decoding of an organism’s genetic information. Translation is a highly regulated process, and this regulation is important for tuning protein levels, controlling the location of protein production and for quality control of both mRNAs and their synthesized proteins. Deregulation of mRNA translation underlies many pathologies, including neurodegenerative diseases and cancer (Bhat et al., 2015; Tahmasebi et al., 2018), highlighting the importance of accurate translational control.

Translation is composed of three phases; initiation, elongation and termination. In eukaryotic cells, translation initiation generally starts with recruitment of the small ribosomal subunit complexed with several initiation factors to the 5’ end of the mRNA. After recruitment, the ribosome scans along the 5’ UTR to identify a translation initiation codon (Aitken and Lorsch, 2012; Brito Querido et al., 2023). During this scanning phase, the helicase eIF4A associates with the small ribosomal subunit and is thought to unfold mRNA structures that might impede ribosome scanning (Andreou and Klostermeier, 2013; Yourik et al., 2017). Upon identification of the start codon, the large ribosomal subunit is recruited and translation elongation ensues. The translation elongation cycle consists of several steps, including: the decoding step, during which an amino-acetylated tRNA binds to the mRNA codon present in the ribosomal A-site; peptide bond formation, during which the nascent polypeptide chain is transferred from the P-site tRNA to the A-site tRNA to extend the nascent chain by one amino acid, and translocation of the mRNA through the ribosome by 3 nucleotides to allow decoding of the next codon (Behrmann et al., 2015; Dever et al., 2018). Upon entry of a stop codon into the ribosome A-site translation is terminated, causing release of the nascent chain and recycling of the ribosomal subunits (Dever and Green, 2012; Lawson et al., 2021).

Translation is regulated both globally and at the level of specific mRNAs. While the initiation step is the dominant point of regulation for controlling protein synthesis rates, the elongation cycle too is under tight control. Translation elongation rates can control expression levels of proteins directly by controlling protein synthesis rates, but also indirectly through control of mRNA stability (Bae and Coller, 2022; Dave et al., 2023; Radhakrishnan and Green, 2016). The rate of translation elongation not only determines protein expression levels, but also protein quality; for example, several studies have found that ribosome translocation speeds affect nascent polypeptide folding (Crombie et al., 1992; Gloge et al., 2014; Liutkute et al., 2020). A pause in translation elongation is also critical for correct targeting of transmembrane proteins to the endoplasmic reticulum (ER) (Collart and Weiss, 2020). Additional elongation pause sequences exist in specific mRNAs, including in the transcription factor Xbp1, which allows expression of Xbp1 under stress conditions (Yanagitani et al., 2011). Viral RNAs also frequently encode translation pause sequences, for example to induce ribosome frame-shifting, which is employed to encode different proteins in a single RNA sequence (Atkins et al., 2016). Thus, regulation of translation elongation rates is critical to control protein levels and function.

Mechanistically, translation elongation rates can be controlled in a variety of different ways. Global elongation rates are controlled by phosphorylation of elongation factor 2 (eEF2) in response to a variety of intracellular and extracellular signals (Dever *et al*., 2018; Proud, 2019). Elongation rates can also be controlled at a gene-specific level. For example, differential codon usage can control elongation rates for a specific mRNA, since the expression levels of tRNAs impact the decoding speed of their cognate codons (Gobet et al., 2020; Hanson and Coller, 2018; Neelagandan et al., 2020). In addition, nascent polypeptides can also affect elongation rates through interactions with the ribosome exit tunnel or by modulating kinetics of peptide bond formation (Collart and Weiss, 2020; Gutierrez et al., 2013). Furthermore, strong RNA structures in the coding sequence of an mRNA are thought to slow down ribosome translocation as well (Wen et al., 2008). Regulatory proteins can also slow down elongation, including the signal recognition particle (SRP) that binds to and pauses ribosomes translating transmembrane and secreted proteins (Halic et al., 2004), and Argonaute proteins complexed with miRNAs (Sako et al., 2023). In addition to physiological regulation, elongation rates can also be altered by damage to ribosomes or mRNAs. For example, oxidation or UV induced damage to mRNAs can cause ribosomes to stall (Snieckute et al., 2023; Yan et al., 2019).

Despite the importance of translation elongation regulation, it has remained surprisingly challenging to study the translation elongation phase. *In vitro* single-molecule FRET and biochemical studies have determined the kinetics of each sub-step in the elongation cycle (Blanchard et al., 2004; Caliskan et al., 2014; Uemura et al., 2010), but don’t capture the complexity or heterogeneity of translation in cells. *In vivo*, several methods have been developed to measure translation elongation, which include methods to measure average, global translation elongation rates (Argüello et al., 2018) and methods to measure genome-wide average decoding time of individual codons (Brar and Weissman, 2015; Ingolia et al., 2009). A major limitation of earlier methods is that they provide only an average elongation rate of a cell, mRNA or codon. More recently, we and others have developed an approach to measure translation dynamics of single mRNAs by fluorescent labeling of the nascent polypeptide using the SunTag (Pichon et al., 2016; Wang et al., 2016; Wu et al., 2016; Yan et al., 2016) or similar antibodies (Morisaki et al., 2016). While these approaches provide single mRNA resolution, a major drawback of these methods is that typical mRNA molecules are translated by many ribosomes simultaneously, obscuring the kinetics of individual ribosomes during elongation. Additionally, elongation rate measurements on single mRNAs are very noisy both due to the low fluorescent signal associated with single translating mRNAs and the stochastic nature of translation. Since almost all our knowledge on *in vivo* translation elongation dynamics comes from averaged measurements over many ribosomes, little is known about the behavior of individual ribosomes during translation elongation.

Here, we develop a method to measure translation elongation rates of single ribosomes with very high precision. We generated circular RNAs that lack in-frame stop codons (called socRNAs; <u>S</u>topless-<u>O</u>RF <u>C</u>ircular <u>RNA</u>s). Either single or multiple ribosomes can be loaded onto socRNAs, allowing analysis of individual ribosomes, but also of functional interplay between ribosomes (Madern et al., 2024). Translation of socRNAs is visualized using the SunTag translation imaging system, allowing us to follow translation of socRNAs by individual ribosomes for hours and to measure their elongation rates over time. We apply the socRNA method to accurately measure pause durations at rare codons and pause-inducing peptide sequences, and we identify a highly non-linear relationship between pause duration and structural stability at RNA structures. In addition, we show that different ribosome-targeting drugs have distinct effects on elongation dynamics, which has important implications for their application. We also identify a novel role for the translation initiation factor eIF4A in stimulating elongation, which was made possible by the ability of socRNAs to experimentally uncouple translation initiation from elongation. Moreover, detailed investigation of single ribosome translocation rates revealed that ribosomes undergo infrequent prolonged pauses and that different ribosomes move at slightly different speeds. Finally, we adapt our socRNA approach to study translation elongation fidelity, which revealed that single ribosomes undergo frameshifting at low, but detectable rates at non-repetitive sequences. Together, our study uncovered detailed translation elongation dynamics of individual ribosomes *in vivo* and provides a powerful, easy-to-use and broadly-applicable new technology to study translation elongation.

## Results

To generate Stopless-ORF circular RNAs (socRNAs) that could be translated in cells, we used the previously developed Tornado system (Litke and Jaffrey, 2019), which is based on a linear precursor RNA that contains two ribozymes which cleave the RNA, followed by ligation of the 5’ and 3’ ends of the excised RNA fragment by RtcB cellular ligase (Figure 1A). To visualize translation of socRNAs, we used the SunTag translation imaging system that we and others have previously developed (Morisaki *et al*., 2016; Pichon *et al*., 2016; Wang *et al*., 2016; Wu *et al*., 2016; Yan *et al*., 2016). In brief, as the ribosome translates the RNA sequence of the socRNA encoding the SunTag peptides (five SunTag peptides are encoded per socRNA, unless stated otherwise), the nascent SunTag peptides emerge from the ribosome and are co-translationally labeled by the SunTag antibody (STAb), which is tagged with GFP and stably expressed at low levels in the cell (Figure 1B). All stop codons were removed from the socRNAs in the SunTag translation reading frame and the socRNA sequence was designed such that it contained a multiple of 3 nucleotides, to ensure that the ribosome remained in the same reading frame upon completing a full circle. We have previously shown that long-term mRNA tracking and signal-to-noise in imaging are both enhanced when the SunTag mRNAs are tethered to the plasma membrane (Yan *et al*., 2016). To ensure similar imaging precision, we also wanted to tether socRNAs to the plasma membrane. However, we tethered linear mRNAs to the membrane through a membrane-anchored protein that binds to the 3’UTR of the mRNA, which is not possible for socRNAs, as they do not contain an untranslated region, so any RNA-membrane tether would be displaced from the RNA by a translating ribosome. We therefore developed an alternative approach in which socRNAs are tethered to the membrane through their nascent chain (Figure 1B). Nascent chain tethering was achieved by introducing the sequence encoding a second, orthogonal epitope tag, the ALFA-tag (Bellec et al., 2023; Gotzke et al., 2019), in the socRNA and by expressing the ALFA-tag nanobody (ALFANb) fused to a membrane anchor in cells (Figures 1B and 1C). Finally, we engineered a doxycycline-inducible promoter to drive expression of the socRNA, so that its expression can be temporally controlled to capture the early phase of socRNA translation.

**Figure 1.**
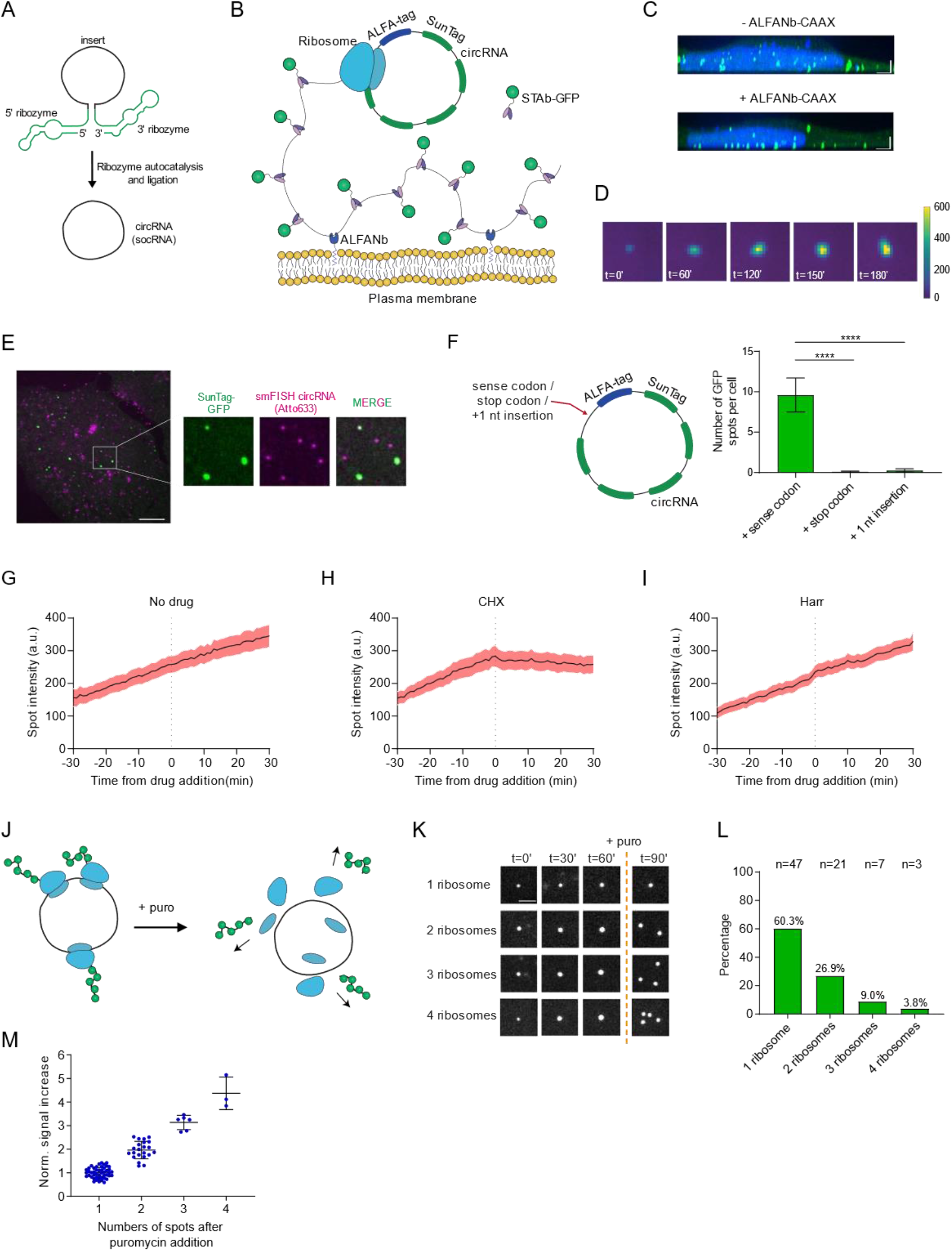
A method for long-term visualization of single translating ribosomes in living cells. **A)** Illustration depicting the generation of socRNAs using the Tornado system. **B)** Schematic of socRNA system. Translation of socRNAs is visualized through binding of GFP-tagged SunTag antibody (STAb-GFP) to nascent SunTag peptides. socRNAs are tethered to the plasma membrane through binding of ALFA-tag peptides to the ALFA-tag nanobody (ALFANb), which is tethered to the plasma membrane. **C)** Representative images of cells in which socRNAs are either freely diffusing through the cytoplasm (top) or tethered to the plasma membrane through the ALFA-tag system (bottom). Translating socRNAs can be observed as green foci and the nucleus is stained by DAPI (blue). **D)** Time-lapse analysis shows that the GFP intensity associated with a single translating socRNA increases over time, indicative of ongoing translation. **E)** Representative image of fixed U2OS cells expressing socRNAs and STAb-GFP. socRNAs are stained by smFISH. Note that only a small subset of socRNAs is undergoing translation. **F)** Insertion of a stop codon or a single nucleotide (which causes a frameshift) into the socRNA eliminates GFP foci formation. Schematic of socRNAs (left) and quantification of the number of GFP foci per cell (right) are shown. **** indicates p<0.0001, t-test. Error bars indicate standard error of the mean. **G-I)** Intensities of translating socRNAs over time are shown for either untreated cells (F), or cells treated with either cycloheximide (CHX) (G) or Harringtonine (Harr) (H). Line indicates the mean and shaded region indicates standard error of the mean. **J)** Schematic showing that socRNAs translated by multiple ribosomes split into multiple GFP foci upon puromycin treatment. **K)** Time-lapse analysis of socRNAs translated by either 1, 2, 3 or 4 ribosomes. Puromycin is added at the indicated time-point. **L)** socRNA-expressing cells were followed by time-lapse microscopy. Puromycin was added during the movie. GFP signal was measured over time until the moment of puromycin addition and socRNAs were grouped based on the number of translating ribosomes associated with each socRNA (assessed as in (K)). **M)** Relationship between number of spots upon puromycin addition and signal increase over time. The rate of GFP increase for socRNAs translated by 1 ribosome was normalized to a value of 1. Error bars indicate standard deviation. Scale bars, 3 μm (C), 10 μm (E), 2 μm (K). The number of experimental repeats and cells analyzed per experiment are listed in Table S1.

We transfected a plasmid encoding the socRNA in STAb-GFP and membrane anchored ALFANb-expressing human U2OS cells and imaged cells by spinning disk confocal microscopy. Time-lapse imaging revealed many individual GFP foci that increased in intensity over time (Figure 1D and Video S1), consistent with translation of socRNAs. To assess whether these GFP foci indeed represent translating socRNAs, we performed a number of control experiments. First, we fixed socRNA expressing cells after time-lapse imaging and labeled individual socRNAs by single molecule FISH (smFISH) (Figure 1E). GFP foci that were increasing in intensity at the moment of fixation generally showed co-localization with socRNAs, while foci that were not increasing in intensity generally did not co-localize with socRNAs, suggesting that the latter group of foci represents protein products for which translation had been aborted (as will be discussed later) and which were released from the ribosome and socRNA template (Figure S1A). As a second set of controls, we inserted a stop codon in the SunTag frame, or added one additional nucleotide to the socRNA, such that the ribosome would change frames and rapidly encounter a stop codon after completing a full circle of translation (Figure 1F). In both cases, no GFP foci could be observed, presumably because five SunTags are not sufficient to generate observable foci, which demonstrates that an ‘infinite circular ORF’ is required for GFP foci formation. Finally, we added the translation elongation inhibitor cycloheximide (CHX) during time-lapse imaging of socRNAs and found that the GFP increase was acutely blocked upon CHX treatment (Figures 1G and 1H), further confirming that the increase in GFP over time was caused by active translation elongation. As expected, the translation inhibitor harringtonine, which blocks ribosomes at the translation initiation codon but doesn’t affect subsequent elongation, did not inhibit GFP increase over time (Figures 1I and S1B). We also checked whether the very long nascent chains formed by continuous translation of socRNAs inhibited translation elongation, but found that it did not inhibit elongation, as translation elongation rates remain constant during prolonged translation elongation (See Figure 5). Thus, we conclude that socRNAs allow long-term measurements of translation elongation rates.

To determine the number of ribosomes translating individual socRNA molecules, we treated cells with the translation inhibitor puromycin, which releases nascent chains from ribosomes. If socRNAs are translated by multiple ribosomes simultaneously, GFP foci should split into multiple smaller foci upon puromycin treatment, as the nascent chains are released from ribosomes and can freely diffuse away from each other (Figure 1J). While some GFP foci remained a single spot upon puromycin treatment, others rapidly split into two or more smaller foci (Figures 1K-1L and Video S2). Foci that split into two smaller spots showed an approximately two-fold higher rate of GFP accumulation than foci that did not split (Figures 1M and S1E), consistent with the notion that these socRNAs were translated by two ribosomes simultaneously. A similar relationship between daughter foci number and the rate of increase in GFP intensity was observed for foci that split into three or four daughter spots (Figure 1M). In contrast, the smFISH foci intensity did not correlate with the number of translating ribosomes per GFP spot, demonstrating that all ribosomes present in individual GFP foci were translating a single socRNA (Figure S1C). In summary, these results show that socRNAs can be translated by one or more ribosomes simultaneously, and that addition of puromycin at the end of an experiment allows a straightforward analysis of the number of ribosomes translating each socRNA.

To further characterize the socRNA system, we next asked how ribosomes are loaded on socRNAs. We considered two possible mechanisms for ribosome loading; first, 43S ribosomes could be slotted directly onto the circular form of the RNA. Alternatively, it is possible that ribosomes are loaded on the linear precursor RNA through conventional 5’ cap-dependent loading, and that these ribosomes are ‘caught’ as the RNA circularizes while they are translating the region of the linear RNA that ends up in the circRNA (Figure S1D). An important hint into the ribosome loading mechanism came from the intensities of GFP ‘daughter’ foci after puromycin-induced splitting, which were almost always exactly equal (Figures S1E-S1G). Equal intensities of daughter foci indicates that all ribosomes translating the same socRNA initiated translation at a similar time, which is consistent with the 5’ loading and capture model. To further confirm the 5’ loading and capture model, introduced an AUG start codon in the 5’UTR of the linear precursor RNA upstream of the socRNA sequence in different reading frames. We reasoned that an AUG start codon in the 5’UTR would only influence the reading frame of ribosomes translating the mature socRNA if ribosomes are loaded through the 5’ loading method, but not if ribosomes are loaded through direct slotting onto the socRNA, as the additional AUG is not present in the mature socRNA. To assess the reading frame that ribosomes are translating, we designed socRNAs encoding both SunTag and ALFA-tag, but in different stopless reading frames, analogous to our previous reading frame reporters on linear mRNAs (Boersma et al., 2019; Lyon et al., 2019). We then expressed these dual-frame socRNAs in a cell line expressing STAb-GFP and ALFANb-HaloTag to visualize translation in both reading frames simultaneously. Introduction of an upstream AUG in the ALFA-tag reading frame strongly increased the relative number of ribosomes translating the ALFA-tag reading frame (Figures S1H-S1J), confirming that ribosomes translating socRNAs initiated on the linear precursor RNA.

Having established socRNAs as a robust and reliable assay to measure translation elongation by single ribosomes, we set out to determine whether socRNAs could be used to precisely measure ribosome translocation dynamics at problematic sequences. We first introduced a known pause sequence from the stress-induced transcription factor Xbp1 into the socRNA (referred to as Xbp1 socRNA) (Figure 2A). The Xbp1 pause sequence blocks elongation at a precisely defined site in the mRNA through interactions of the polypeptide with the ribosome exit tunnel (Shanmuganathan et al., 2019). For control socRNAs lacking the Xbp1 pause site, we calculated that ribosomes translate at a rate of 2.5 codons/s (See Methods), a rate that is similar to our previous elongation rate measurements (∼3 codons/s) on linear mRNAs in the same cells (Yan *et al*., 2016). Comparing translation elongation rates for control and Xbp1 socRNAs translated by a single ribosome revealed a substantially slower average translation elongation for Xbp1 socRNAs (Figure 2B). Based on the differences in the rate of GFP intensity increase we could calculate that the pause duration to be 106 sec on the Xbp1 pause sequence (Figure 2C). Pausing on the different pause sequences derived from the human cytomegalovirus (hCMV) gp48 (Bhushan et al., 2010) and fungal arginine attenuator peptide (AAP) (Wei et al., 2012) could also be precisely measured to be 41 and 42 s, respectively. These results show that socRNAs accurately recapitulate known pausing sequences, and enable precise quantitative assessment of pause duration. In addition to measurements of translation elongation rates, the socRNA assay also uniquely allows measurements of ribosome processivity (defined here as the total number of codons translated until translation is aborted). Ribosome processivity could be affected by a number of different processes, including 1) translation termination on sense codons, 2) ribosome recycling in the absence of termination - such as through activation of quality control mechanism (Joazeiro, 2017), 3) ribosome frameshifting followed by termination on a stop codon in the alternative reading frame, or 4) the decay of socRNAs (see Discussion section). Analysis of ribosome processivity on control socRNAs showed that ribosomes are highly processive, translating on average ∼20,000 codons before aborting translation (Figures 2D and 2E), which is ∼60-fold more than the length of an average mRNA in human cells. Interestingly, for ribosomes translating Xbp1 socRNAs, the total number of translated codons is reduced by 1.4-fold (Figure 2E), showing that a strong pause sequence can reduce ribosome processivity. Importantly, individual ribosomes still translate the pause sequences >30 times, indicating that ribosomes remain attached to the mRNA and are capable of resuming translation even after 100 seconds long pauses. Furthermore, these results show that socRNAs provide an incredibly sensitive readout of ribosome processivity, allowing detection of a <1% chance of aborting translation when ribosomes encounter a problematic sequence.

**Figure 2.**
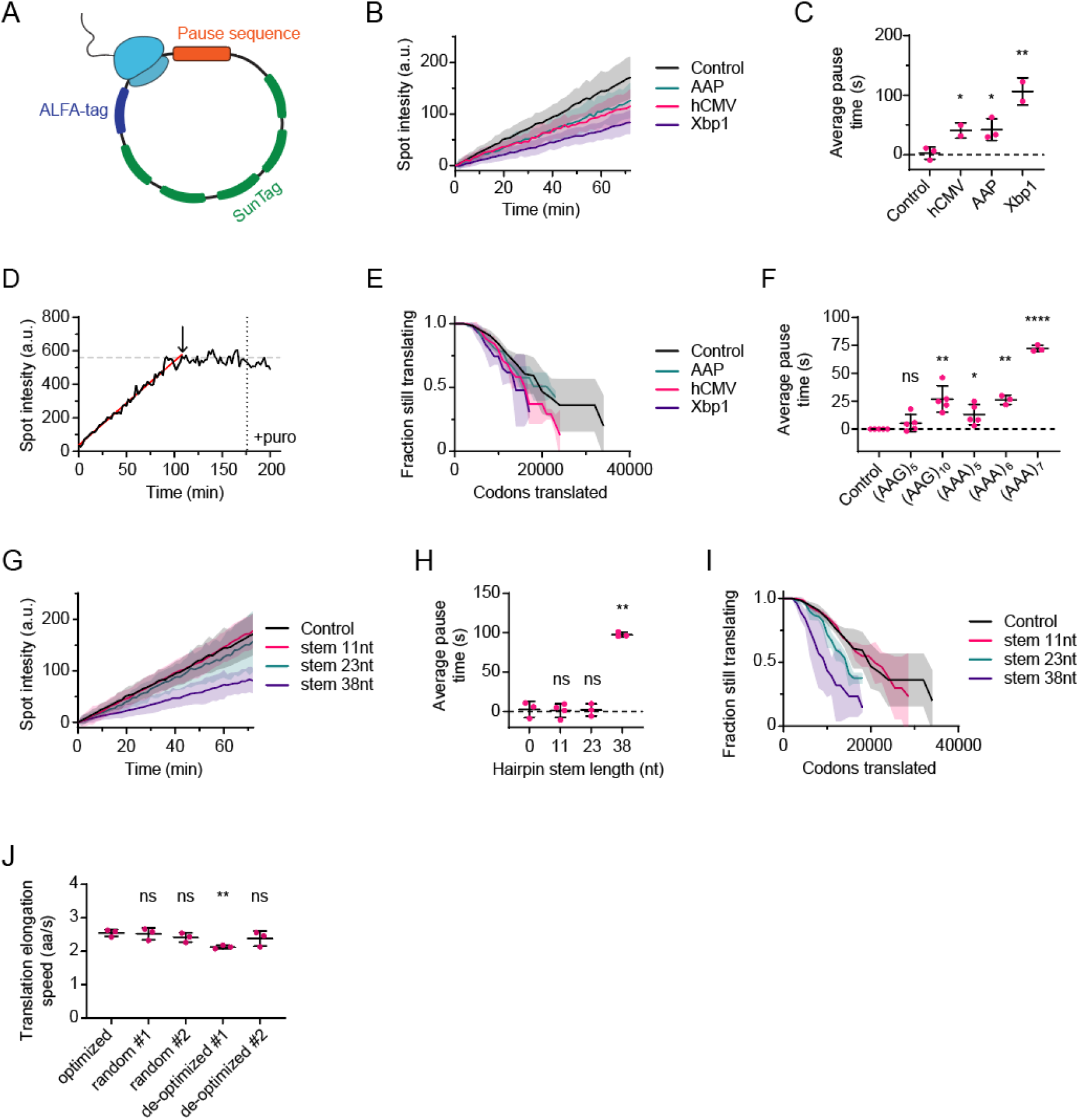
Precise measurements of ribosome pausing and processivity. **A)** Schematic of socRNA with an introduced translation pause sequence. **B-J)** U2OS cells stably expressing STAb-GFP, ALFANb-CAAX, and tetR were transfected with indicated socRNAs and imaged by time-lapse microscopy. **B, G)** SocRNA GFP foci intensity was measured over time. The intensities at the start of the measurement were set to 0. **C, F, H)** Pause durations for each time a ribosome encounters the pause sequence was calculated (See Methods). **D)** Representative GFP intensity time trace of a socRNA showing abortive translation before puromycin addition. Dashed vertical line indicates moment of puromycin addition. The moment when the GFP intensity stopped increasing was determined to calculate the number of codons translated by individual ribosomes on socRNAs. Red line indicates the increasing phase of the GFP spot intensity. The arrow indicates the moment of translation aborting. **E, I)** Kaplan-Meyer survival curve showing the total number of codons translated by ribosomes before aborting translation. **J)** Translation elongation speed was measured for 5 socRNAs encoding the same amino acid sequence but differing in their codon optimality. Codon Adaptation Index (CAI) for respective socRNAs, from left to right: 0.84, 0.67, 0.66, 0.49 and 0.51, respectively. Lines in (B, E, G, I) indicate mean values, shaded regions indicate standard deviation. *, ** and **** indicate p<0.05, 0.01 and 0.0001 respectively (t-test). Error bars indicate standard deviation from independent experiments. The number of experimental repeats and cells analyzed per experiment are listed in Table S1.

We also examined ribosome pausing on polylysine stretches, which pause ribosomes through their interaction with the ribosome exit tunnel due to their high positive charge (Arthur et al., 2015; Lu and Deutsch, 2008). socRNAs allow quantitative and very sensitive measurements of pausing on different lengths of polylysine stretches. In addition, socRNAs allow measurements of *single* ribosomes translating polylysine stretches, which is important as we found that additional ribosomes strongly suppress pausing on polylysine stretches (Madern et al., 2024). We found a pause duration of 5 s on 5 consecutive AAG-encoded lysine residues (Figure 2F). A doubling of the lysine stretch to ten consecutive lysine residues caused ribosomes to pause >5-fold longer (27 s) (Figure 2F), indicating that an increased polylysine stretch synergistically increases pause duration, possibly through increased avidity in the polylysine-ribosome exit tunnel interaction. Consistent with previous reports, we find that a stretch of 5 AAA lysine codons resulted in a somewhat stronger pause than 5 AAG lysine codons (5 vs 13 sec), likely due to a unique structure formed by consecutive AAA codons that slows down decoding (Chandrasekaran et al., 2019; Tesina et al., 2020). Surprisingly, increasing the AAA codon stretch by just 2 codons, from 5 to 7 consecutive AAA codons, dramatically increased the pause duration from 13 to 72 sec (Figure 2F), a far stronger synergistic effect than was observed with AAG lysine codons, suggesting that the inhibitory effect of an adenosine stretch scales exponentially with the length of the nucleotide sequence. Together, these results show that the sensitive and highly quantitative nature of the socRNA assay can provide improved understanding of polylysine translation, as well as other repetitive sequences that impede translation.

Next, we examined translation elongation kinetics of structured RNAs. GC-rich hairpins were introduced into the socRNA with variable stem lengths (Figure 2G), and pause durations were calculated for socRNAs translated by a single ribosome (Figure 2H). No pause was detected for hairpins with stems of 11 or 23 nt, demonstrating that ribosomes are incredibly efficient in unfolding RNA structure during translation elongation. Less than 0.5% of human transcripts contain predicted RNA structures with a folding energy that is higher than the 23 nt hairpin (Rouse et al., 2022), indicating that ribosomes can efficiently translate the large majority of the transcriptome without pausing due to RNA structure. However, hairpins with 38 nt stems did slow down elongation substantially, inducing average pauses of 98 s (Figure 2H), showing that there is a limit to the unfolding capacity of ribosomes. Examination of the total translated codons revealed little effect on ribosome processivity of the 11 and 23 nt stem structures, but a 2-fold reduction for the 38 nt stem (Figure 2I). Nonetheless, ribosomes still unfolded the 38 nt stem on average 31 times before aborting translation, further confirming the highly processive nature of translation elongation and the potency of the ribosome in unfolding RNA structures during elongation.

In addition to stalling peptides and RNA structures, codon usage is also thought to alter translation elongation rates (Hanson and Coller, 2018). Previous work has shown that ‘non-optimal codons’, are decoded more slowly than ‘optimal’ codons. To test the effect of codon optimality on translation elongation rates, we introduced synonymous mutations into socRNAs for 89% of all codons (note that a small region of the socRNAs cannot be mutated, as it is essential for RNA circularization). We either used the most optimal, random, or non-optimal codons (see Methods), according to the codon adaptation index, while maintaining identical amino acid sequences for all socRNAs (Figure 2J). Somewhat surprisingly, we found that even this very extreme (de-)optimization of codon usage had very limited effects on translation elongation rates, with the optimized socRNA showing an almost identical translation elongation rate to the codon-randomized reporters (2.54 vs. 2.51 and 2.41 codons/s), and the two de-optimized socRNAs showing only a 6.5% and 16.4% reduction in elongation rates compared to the fully optimized sequence (2.54 vs 2.38 and 2.13 codons/s). Thus, we conclude that codon optimality does not have a major effect on average translation elongation rates, despite having a substantial impact on gene expression (Presnyak et al., 2015; Wu et al., 2019).

The ribosome is a frequent target of small molecules produced by a variety of microorganisms. Ribosome targeting molecules are widely used in biomedical research to experimentally modulate translation elongation dynamics and also as therapeutic agents in the clinic (Carocci and Yang, 2016; Lin et al., 2018; Panwar et al., 2020). While the precise mechanism of action has been resolved from a structural perspective for a small number of ribosome targeting drugs (i.e., identification of the binding site on ribosomes (Garreau de Loubresse et al., 2014; Schneider-Poetsch et al., 2010)), the effects of these drugs on translation elongation dynamics *in vivo* are poorly understood. Understanding how such drugs affect translation elongation dynamics is often critical for correct interpretation of experiments and for optimal clinical application. For example, intermediate doses of a ribosome targeting drug may appear to slow down translation elongation in a bulk experiment, but an apparent slowdown could be caused either by slowing down all ribosomes equally over time, or by complete immobilization of a subset of ribosomes, while leaving other ribosomes unchanged. Indeed, at intermediate concentrations the translation elongation inhibitor anisomycin induces widespread ribosome collisions (Juszkiewicz et al., 2018; Wu et al., 2020), indicative of arrest of a subset of ribosomes. To directly assess how different ribosome targeting drugs affect translation elongation dynamics, we treated cells expressing socRNAs with different elongation inhibitors – CHX, anisomycin, and narciclasine – and measured their dissociation kinetics from the ribosome *in vivo*. At high concentration, all three drugs completely inhibited translation elongation, as expected (Figure S2A). Treatment of cells with high doses of drugs, followed by drug washout (Figure 3A) revealed that CHX dissociates very rapidly from arrested ribosomes (Figures 3B and 3F), whereas anisomycin and narciclasine induced very long-lived ribosome stalls (>10 minutes) (Figures 3C, 3D, and 3F). These results reveal the *in vivo* dynamics of commonly-used ribosome targeting drugs, and provide a powerful assay to measure the dynamics of other drugs on ribosome translocation kinetics.

**Figure 3.**
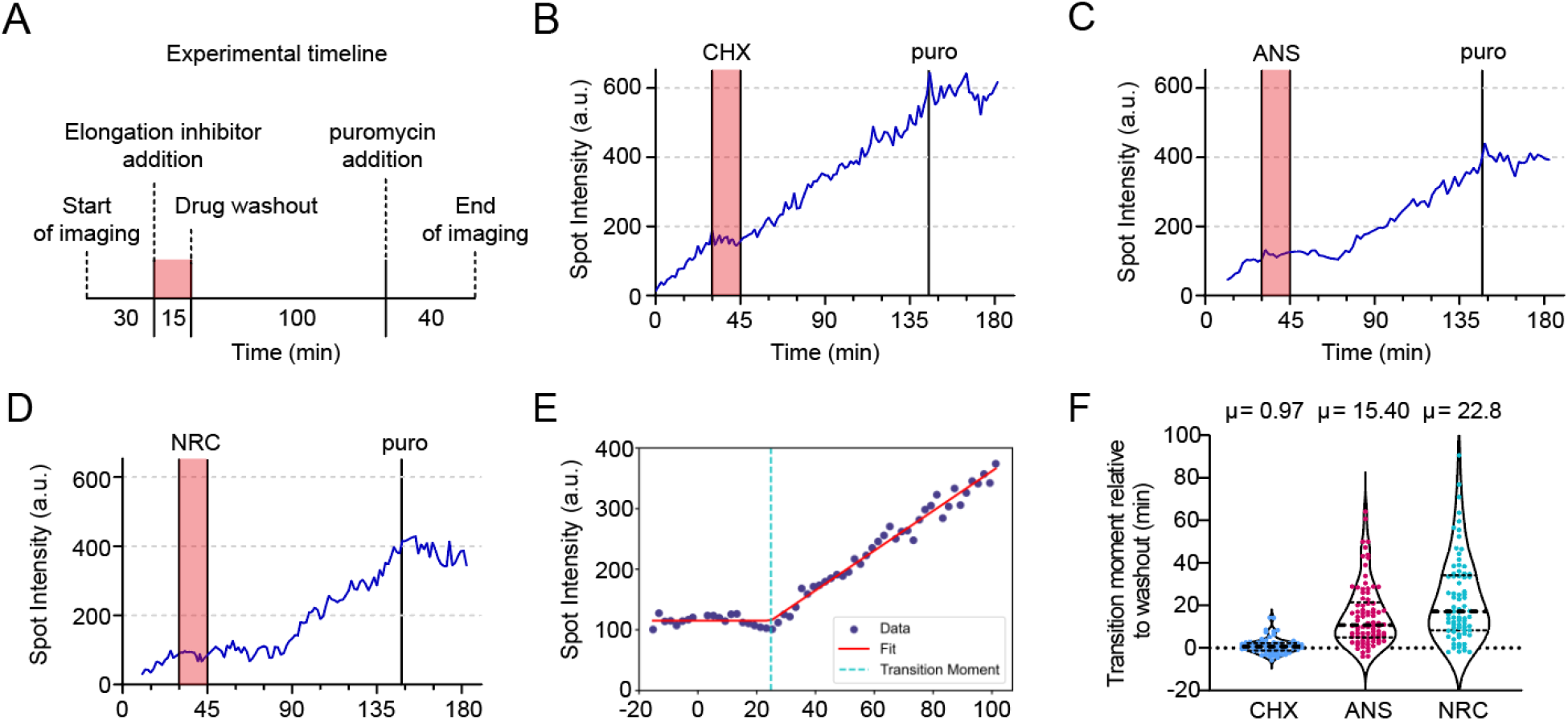
Dissecting dynamics of ribosome targeting drugs. **A)** Overview of the experimental setup used in (B-F). **B-D)** Representative intensity-time traces of single translating socRNAs treated with indicated ribosome-targeting drugs. **E)** Representative example of data fitting approach to identify the moment that translation resumes after translation pausing induced by ribosome targeting drugs. **F)** The time from drug removal to resumption of translation is shown for three different ribosome targeting drugs. Each spot represents a single translated socRNA tracked over time. The thick dashed line indicates the median, thin lines indicate 25th and 75th percentile. The number of experimental repeats and cells analyzed per experiment are listed in Table S1.

Another unique aspect of the socRNA translation elongation assay is that elongation can be measured even after global translation initiation shutdown in the cell, as socRNAs do not require continued translation initiation for elongation measurements (as long as initiation occurred before global initiation shutdown). Many biological processes are known to cause global shutdown of translation initiation, including widespread protein misfolding in the ER, nutrient starvation and viral infection (Reid and Nicchitta, 2015; Shu et al., 2020; Walsh and Mohr, 2011). In many cases, a few mRNAs are thought to escape the global initiation shutdown, for example innate immune genes during viral infection (Rozman et al., 2023), but it has been difficult to assess translation elongation dynamics under such conditions with existing assays. We first asked whether global inhibition of translation affects elongation rates, for example by increasing the availability of charged tRNAs or elongation factors. Expression of socRNA transcription was induced and after 135 min cells were treated with harringtonine to shut down global translation, while allowing continued translation on socRNAs (see Figure 1I). Translation elongation rates were measured on socRNAs before and after harringtonine addition for the same socRNAs, which revealed that translation elongation occurs at very similar rates before and after global suppression of translation (Figures 4A and 4B). These results show that translation elongation can be assessed during global translation initiation inhibition using socRNAs and indicate that in unperturbed, high nutrient conditions, cellular resources are not limited for translation elongation.

**Figure 4.**
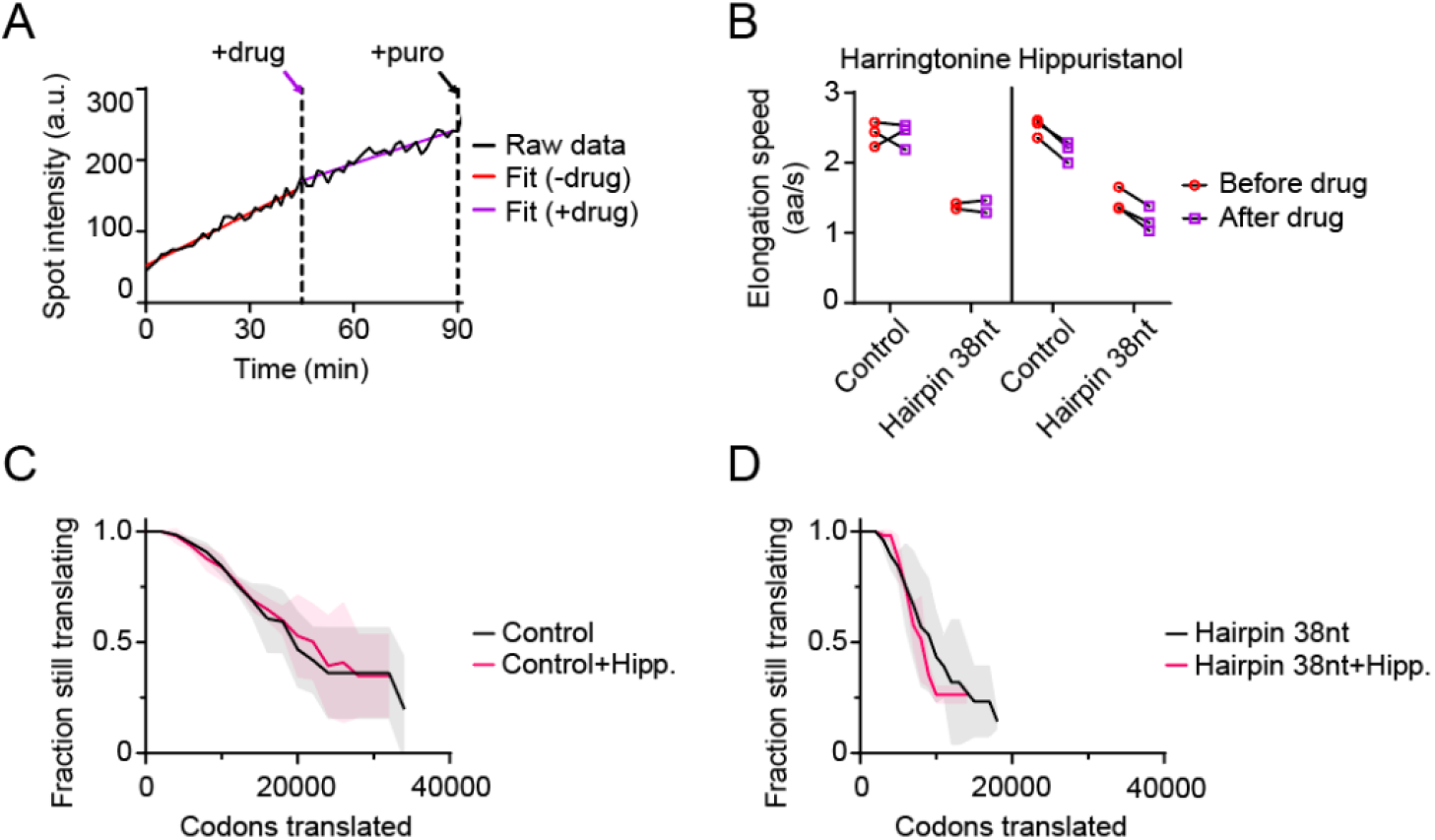
eIF4A promotes translation elongation. **A-D)** U2OS cells stably expressing STAb-GFP, ALFANb-CAAX, and tetR were transfected with indicated socRNAs and imaged by time-lapse microscopy. Cells were treated with either the early elongation inhibitor harringtonine or the eIF4A inhibitor hippuristanol, as indicated. At the end of the experiment, cells were treated with puromycin to assess the number of ribosomes translating each socRNA. **A)** Representative example intensity time trace of a single socRNA (black line). Moment of drug addition is indicated by dashed vertical lines. Intensity time trace was split into two sections (before and after drug addition) and the best linear fit for section part was determined (red line before drug addition and purple line after drug addition). **B)** Elongation rates were calculated before (red circle) and after drug (purple square) addition for the same socRNAs for indicated socRNAs and drug treatments. **C-D)** Kaplan-Meyer survival curve of indicated socRNAs and drug treatments showing the total number of *codons* translated by ribosomes before aborting translation. Lines indicate mean values, shaded regions indicated standard deviation. The number of experimental repeats and cells analyzed per experiment are listed in Table S1.

Next, we tested the role of the helicase eIF4A, a well-established translation factor that is part of the eIF4F translation initiation complex (Aitken and Lorsch, 2012), in translation elongation. While it is known that eIF4A contributes to translation *initiation* by unwinding RNA during ribosome recruitment and/or ribosome scanning along the 5’UTR (Yourik *et al*., 2017), it is unknown if eIF4A is also involved in translation *elongation*. Because translation initiation is completely shut down upon inhibition of eIF4A, studying potential roles of eIF4A in elongation has been challenging. We reasoned that socRNAs would allow uncoupling of translation initiation and elongation, and thus allow assessment of a role of eIF4A in translation elongation. To this end, we induced expression of socRNAs and added the eIF4A inhibitor hippuristanol (Bordeleau et al., 2006) to cells after 135 min, when initiation had occurred on many socRNAs. Intriguingly, inhibition of eIF4A reduced translation elongation rates by 13%, indicating that eIF4A is important for efficient translation elongation, as well as for initiation (Figure 4B). Considering the role of eIF4A in resolving RNA structure during scanning, we also examined whether the slowdown of translation elongation would be exacerbated on RNA sequences with strong RNA structures. We therefore tested the effects of hippuristanol treatment on translation elongation rates of the socRNA containing the 38nt hairpin structure (Figure 4B). Somewhat surprisingly, introduction of the RNA structure did not further increase the dependency on eIF4A for translation elongation, suggesting that the role of eIF4A in elongation may be distinct from RNA structure unfolding. Finally, we examined whether eIF4A inhibition also affected ribosome processivity, but did not observe a change in processivity upon eIF4A inhibition (Figures 4C and 4D). Together, these results show that eIF4A plays a previously unappreciated role in translation elongation, but is not essential to unfold strong secondary structures during elongation.

A number of studies have shown that ribosomes can vary in composition and that such compositional heterogeneity may be functionally relevant for different aspects of translation (Gay et al., 2022; Genuth and Barna, 2018). The ability to study translation kinetics of individual ribosomes provides a critically-needed tool to assess functional consequences of ribosome heterogeneity. To determine if different ribosomes translate RNAs at distinct speeds, we further improved the accuracy of our elongation speed measurements by correcting for minor movements of GFP foci in the z-direction and for complex photobleaching effects (see Methods). When measuring intensities over time of GFP foci representing single ribosomes translating a socRNA, we found that rates of GFP increase varied considerably between different ribosomes, demonstrating that different ribosomes indeed move at distinct speeds (Figure 5A). To control for technical noise in these measurements, we examined intensity time traces of GFP foci that did not increase in intensity over time (‘plateau traces’), which have similar technical noise. To compare plateau traces with ‘increasing traces’, we transformed the slope of plateau traces with a fixed value, equal to the average slope of increasing traces (Figure 5B, compare black and red distributions). While plateau traces did show some heterogeneity in their slopes as well, the heterogeneity was substantially smaller than that of increasing traces (Figure 5C), demonstrating that technical noise cannot explain the heterogeneity in translation elongation rates, and confirm that different ribosomes move at different speeds. Using the plateau traces as technical noise, we could estimate the actual heterogeneity in translation elongation rates of different ribosomes to be ∼19 % of the mean elongation rate (Figure 5C). Heterogeneity in translation elongation rates was further confirmed using an independent theoretical approach (Figures 5D and S3A-S3E, see Methods).

**Figure 5.**
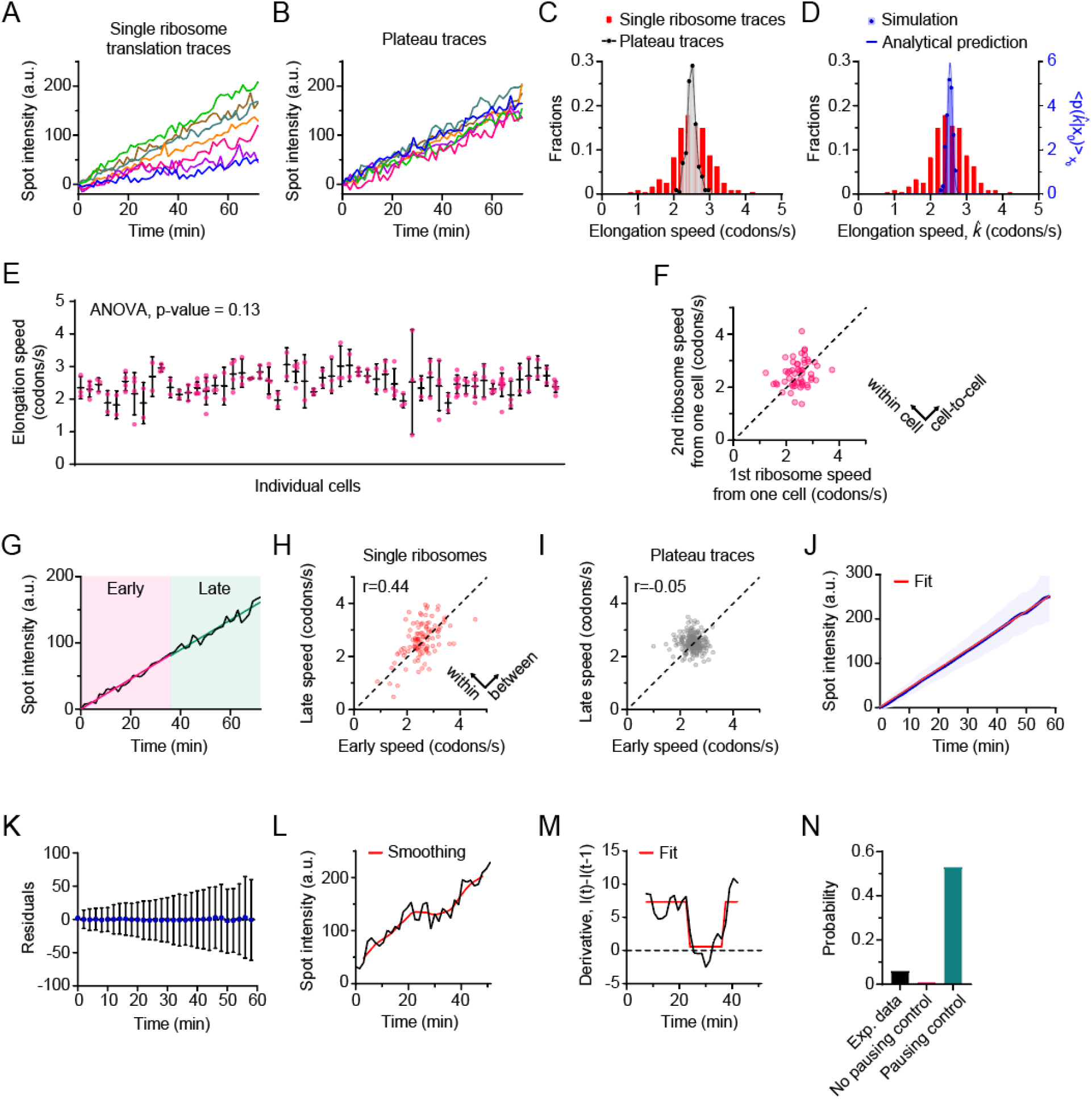
Heterogeneity in single ribosome elongation speeds. **A-O)** U2OS cells stably expressing STAb-GFP, ALFANb-CAAX, and tetR were transfected with indicated socRNAs and imaged by time-lapse microscopy. **A)** Representative intensity time traces of socRNAs translated by individual ribosomes. The intensities at the start of the measurement were set to 0. **B)** Representative control intensity time traces, which indicate the technical noise in intensity time traces. Intensities of GFP foci that were not increasing in intensity (‘plateau traces’) were measured over time and transformed using the mean slope of the intensity time traces of single ribosomes translating socRNAs (See Methods). The intensities at the start of the measurement were set to 0. **C)** Calculated translation elongation rates of all individual ribosomes translating socRNAs (red bars) or transformed control traces (gray bar, plateau traces). **D)** The distribution of elongation speed 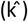 with the distribution 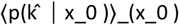 in blue, x_0 represents the estimated starting length of the polypeptide chain (See Methods for details). Predictions from the model are significantly narrower compared to the experimental data, indicative of heterogeneity in the translation elongation rates k between different ribosomes. **E)** Average translation elongation speed on individual socRNAs in different cells. Red dots represent individual socRNAs translated by one ribosome. All magenta dots in each vertical row are from the same cell. Horizontal black lines represent mean and errors bars represent standard deviations. ANOVA statistical test indicates that average elongation speeds in different cells are not statistically different. **F)** Elongation speed of two randomly selected ribosomes translating different socRNAs within the same cell are plotted (the speed of one ribosome is plotted on the x-axis, the other on the y-axis). Note that there is little correlation between elongation speeds of ribosomes within the same cell. Spread of points perpendicular to the diagonal dashed line corresponds to the difference in elongation speeds within the same cell. **G-I)** Slope of the first half (early speed) and second half (late speed) of intensity time traces was determined using a linear fit. **G)** Representative intensity time trace and fitting strategy. **H)** Relationship between the elongation rate of the first half and second half of intensity time traces is shown. Spread over the axis of the dashed line (y = x) indicates heterogeneity in elongation rates between different translating ribosomes. In contrast, spread over the orthogonal axis (y = -x) suggest that ribosomes speed up or slow down during translation of a single socRNA (within trace elongation speed heterogeneity). **I)** Relationship between the slope of the first half and second half of control intensity time traces is shown. **J)** Average GFP intensity over time for all socRNAs combined (black line) and linear fit (red line). Shaded areas around black line represents the standard deviation. The intensities at the start of measurement were set to 0. **K)** Deviation of the experimental data in (J) from the linear fit over time. Note that the data does not deviate more from the linear fit at later time points, demonstrating that ribosomes don’t slow down during socRNA translation over time. **L-N)** Identification of pauses in single ribosome intensity time traces. **L)** Representative raw intensity time trace (black line) and smoothed data (red line, see Methods). **M)** Derivative of the smoothed example trace in (L) (Black line). Red line shows Hidden Markov Modeling to identify translation pauses (plateau’s with a derivative of around 0). **N)** Probability of identifying a pause in intensity time traces of socRNAs using the approach shown in (L,M). As controls, we used transformed plateau traces with/without pause (see Methods). The number of experimental repeats and cells analyzed per experiment are listed in Table S1.

The observed heterogeneity in translation elongation rates could be explained by a number of differences, including: 1) Intrinsic differences in ribosome translocation rates, for example due to differences in ribosome protein composition or rRNA modification, or due to damage to ribosomes, 2) differences in socRNAs, for example due to differential nucleotide sequences, 3) cell-to-cell heterogeneity in translation elongation rates or 4) differences in sub-cellular localization, for example due to attachment to the ER. We first asked whether cell-to-cell heterogeneity in translation elongation rates could explain ribosome elongation rate heterogeneity. When comparing single ribosome elongation rates in the same cell and between different cells, we did not observe a statistically different elongation rate in different cells (Figure 5E), arguing against cell-to-cell heterogeneity as a major driver of ribosome elongation rate heterogeneity. More in depth analysis revealed that cell-to-cell heterogeneity could account for only 22% of the total observed heterogeneity in translation elongation rates (Figure 5F) (See Methods). We next asked whether differences in STAb-GFP expression levels between cells could explain different observed elongation rates, but found no significant correlation between STAb-GFP expression and translation elongation rate (Figure S4A). Additionally, we examined socRNA mobility as a proxy for organelle/membrane association (e.g., ER-localized translation), but found no significant correlation between socRNA mobility and translation elongation rate (Figure S4B). Next, we asked whether rare, stochastic pauses in translation elongation could explain the apparent elongation heterogeneity between ribosomes; if two ribosomes translate at the same rate, but one undergoes a prolonged pause, it would appear to have a slightly slower average translation rate. To address this, we compared elongation rates in the first half and second half of the time traces (Figure 5G) and found a significant correlation between these different time windows (Figures 5H and 5I), indicating that ribosomes maintained a constant speed over a period of tens of minutes, demonstrating that the observed elongation rate heterogeneity was not due to stochastic pausing of ribosomes. We also found that ribosomes do not slow down after a prolonged time of socRNA translation (either because of the large socRNA nascent chain, or because ribosomes become ‘tired’) (Figures 5J and 5K), indicating that elongation rate heterogeneity is not caused by a different total duration of translation. Finally, we examined whether heterogeneity in socRNA sequence, for example through errors in socRNA transcription, could explain elongation rate heterogeneity. We performed sequencing on socRNAs purified from cells, but found no evidence for socRNA sequence heterogeneity (Figure S5). Together, these findings are most consistent with a model in which intrinsic ribosome heterogeneity explains the observed elongation speed heterogeneity (See Discussion).

While stochastic pauses could not (fully) explain the observed translation elongation heterogeneity, we did nonetheless observe occasional pauses in single ribosome intensity time traces (Figure 5L). To systematically quantify pauses in single ribosome intensity time traces, we developed a computational pipeline based on Hidden Markov Modeling (Figure 5M). As a negative control, we used transformed plateau traces (Figure 5C) to account for false positive pause calling due to technical noise. As a positive control, we generated artificial pauses within control traces which were created by transforming plateau traces with a constant positive slope and by introduction of a pause in the middle of the trace to assess pause detection efficiency in our analysis. Using a stringent detection threshold (minimal pause duration 3 min) that resulted in pause calling in less than 1% of control plateau traces, we found that 6% of single ribosome traces showed detectable pausing (Figure 5N). Based on these data we calculated a pause frequency of ∼1/300,000 translated codons, with an average pause duration of ∼11 min. Shorter pauses may occur more frequently, but our assay did not allow identification of brief pauses due to technical limitations. While the observed pause frequency may seem low, extrapolating these results to a typical human mRNA with an CDS length of 325 codons with an initiation rate of 2 min^-1^ (Li and Buck, 2021; Yan *et al*., 2016) reveals that 2.4% of the mRNA molecules could contain a paused ribosome at any given time if these pauses are not resolved. Thus, we conclude that long ribosome pauses are relatively rare, but may impact translational output if left unresolved.

In addition to very sensitive translation elongation rate measurements, we wondered whether socRNAs could also be used for very accurate measurements of translation fidelity. Ribosome frameshifting commonly occurs during translation of viral RNAs in a tightly regulated process to enhance the coding potential of small viral genomes. Such frameshifts are generally induced by a ‘slippery sequence’ (i.e., a repetitive nucleotide sequence) followed by a strong ribosome pause sequence. In addition to regulated frameshifting, frameshifting could potentially also occur as an error in translation, which could lead to synthesis of toxic out-of-frame polypeptides. Frameshifting was shown to occur on poly-adenosine stretches (Arthur *et al*., 2015), but very little is known about the prevalence of frameshifting on non-repetitive RNA sequence, likely because the frequency is below the detection threshold of most assays. SocRNAs may present an opportunity for extremely sensitive measurements of ribosome frameshifting. To measure frameshifting on socRNAs, we generated a dual-color translation reading frame reporter, by inserting one SunTag peptide in one reading frame and an ALFA-tag peptide in one of the two alternative reading frames (Figure 6A). This approach is conceptually similar to imaging-based reading frame reporters we and others have developed previously for linear RNAs (Boersma *et al*., 2019; Lyon *et al*., 2019). All stop codons were removed from both reading frames in our dual-frame socRNAs and an AUG start codon was introduced in the ALFA-tag reading frame, ensuring that most ribosomes initiated translation in the ALFA-tag reading frame (which we refer to as frame 0). In this reporter, production of individual polypeptides containing both ALFA-tag and SunTag peptides is used to assess ribosome frameshifting. We analyzed dual-frame socRNAs by time-lapse microscopy first for socRNAs in which the SunTag was positioned in the +1 frame (Figures 6B-6C and Video S3). As expected, for most socRNAs translation initially occurred in the ALFA-tag reading frame. In 21% of socRNAs ALFA-tag positive foci showed subsequent accumulation of SunTag foci that co-localized with ALFA-tag foci. Importantly, when SunTag signal appeared on ALFA-tag foci, ALFA-tag fluorescence no longer increased (Figures 6C and S6A), consistent with a single ribosome that has undergone frameshifting. Moreover, treatment with puromycin did not result in splitting of ALFA-tag and SunTag signals, excluding the possibility that the SunTag and ALFA-tag translation was performed by two different ribosomes. These results show that socRNAs provide a direct and sensitive readout for ribosome frameshifting and show that ribosome frameshifting does occur on non-repetitive RNA sequences.

**Figure 6.**
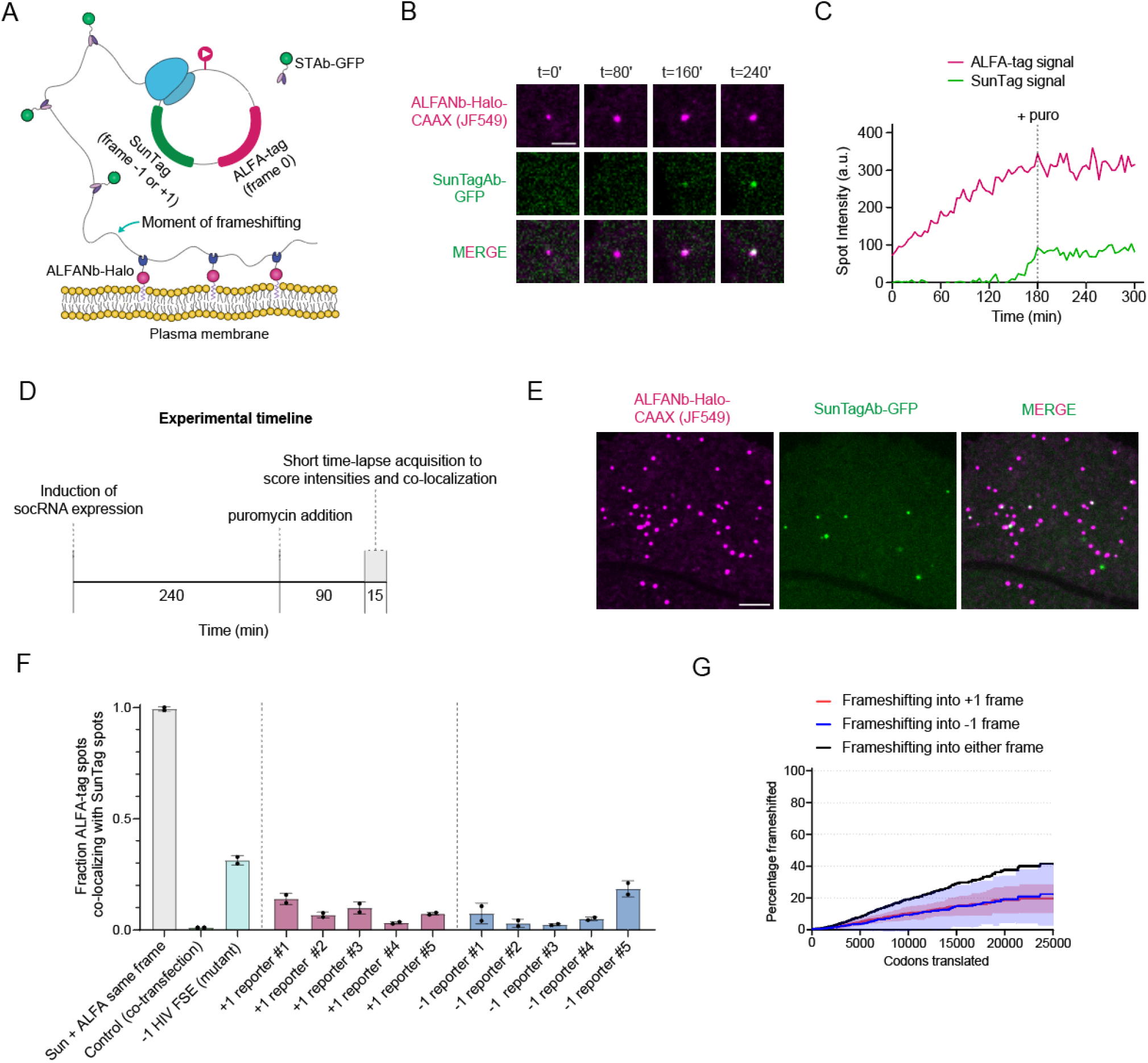
Ultra-sensitive measurements of ribosome frameshifting. **A)** Schematic of ribosome frameshifting assay using socRNAs. socRNAs encode 1 copy of the ALFA-tag peptide and 1 copy of the SunTag peptide in an alternative reading frame, and contain an AUG in the ALFA-tag reading frame (referred to as frameshift socRNA). STAb-GFP is expressed as a cytoplasmic protein, while the ALFA-tag nanobody is attached to a HaloTag (labeled with JF549) and is fused to a membrane anchor (CAAX sequence). Production of a nascent chain results in local accumulation of membrane-bound ALFA-tag-HaloTag nanobodies, which can be observed as a fluorescent puncta. **B-C)** U2OS cells expressing the components described in (A) were followed by time-lapse microscopy and intensity of fluorescent foci was measured for both GFP and HaloTag. Representative images (B) and corresponding intensity time trace (C) of a ribosome frameshifting event into the +1 frame are shown. Dashed vertical line in (C) indicates moment of puromycin addition, which was added to ensure that dual color foci did not reflect a socRNA translated by two different ribosomes in different reading frames (in which cases the two colors should split upon puromycin addition). **D-G)** A snapshot assay was developed to assess frameshifting rates for multiple socRNAs with increased throughput. D) Experimental timeline for snapshot assay to assess frame-shifting with increased throughput. E) Representative image of cell expressing the components described in (A) transfected with socRNA to assess frameshifting into the +1 frame. **F)** Cells stably expressing STAb-GFP and ALFANb-Halo-CAAX were transfected with either one of three control reporters; a socRNA encoding SunTag and ALFA-tag in the same reading frame (green bar, left), two different socRNAs, one encoding the SunTag and the other the ALFA-tag (green bar, middle) or a frameshift socRNA also encoding a weak frameshifting inducing sequence from HIV (green bar, right). In addition, five different frameshift socRNAs were tested, each with randomized nucleotide sequences (but with constant amino acids sequences) in which the SunTag sequence is placed in the +1 frame relative to the AUG sequence and ALFA-tag sequence (magenta bars). Blue bars represent five different reporters with similar design except the SunTag sequences are encoded in the -1 frame. The fraction of ALFA-tag foci that is positive for SunTag signal after puromycin addition (representing frameshifted translation events) is shown for each reporter. Error bars indicate standard deviations. **G)** Kaplan-Meyer survival curve for indicated socRNAs showing the total number of codons translated by ribosomes before frameshifting occurs (See Methods section). Plotted are the average frameshifting rates of all +1 and -1 frame reporters (red line and blue line, respectively) and the sum of both lines (black line), which reflects the total frameshifting rate. Scale bars, 2 μm (B), 5μm (E). The number of experimental repeats and cells analyzed per experiment are listed in Table S1.

To quantitatively assess frameshifting frequencies on non-repetitive sequences, we adapted our assay for increased throughput. We made use of the fact that ribosome frameshifting of dual-frame socRNAs results in dual color (ALFA-tag and SunTag) positive polypeptides. We therefore developed a snapshot assay to measure frameshifting by scoring the fraction of polypeptides that contained both SunTag and ALFA-tag peptides (Figures 6D-6F and S6B-S6C). The single time-point assay revealed a similar frameshifting rate as the live-cell assay (14% vs 21% of ribosomes frameshift during socRNA translation, respectively), and introduction of a weak frameshifting sequence (a mutant variant of the HIV -1 programmed frameshift element (Mouzakis et al., 2013)) significantly increased the frameshifting sequence, together confirming the validity of the snapshot assay. Based on the total number of codons translated and the number of socRNAs that produced frameshifted polypeptides, we could calculate a frameshifting frequency of 1 per ∼42,000 codons translated for our control socRNA (Figure 6G and S6E). To exclude that we had unintentionally introduced a specific sequence that induces frameshifting into the socRNA, we generated four additional +1 dual-frame socRNAs by scrambling the nucleotide sequence of the dual-frame socRNA without altering the amino acid sequence. All four additional socRNAs show frameshifting at similar frequencies (Figures 6F and S6B). We similarly generated five -1 dual-frame socRNAs in which the nucleotide sequence was differentially randomized. All five -1 dual-frame socRNAs showed substantial frameshifting signal as well, at similar frequencies as the +1 socRNAs (Figures 6F and S6C). In summary, these results show that dual-frame socRNAs represent extremely sensitive sensors for ribosome frameshifting, and reveal that ribosome frameshifting occurs at frequencies of around 1 per 42,000 codons on ‘normal’ (i.e., non-repetitive) sequences (Figures 6F and S6E). While this frequency may appear low, at this frequency 0.8% of ribosomes translating an mRNA of 325 codons (median coding sequence of the human transcriptome) will frameshift.

## Discussion

In this study we develop socRNAs (Stopless-ORF circular RNAs) to measure translation elongation with very high precision by tracking single translating ribosomes for hours. Several unique aspects of the socRNA assay make socRNAs uniquely suited to study translation elongation; first, the ability to track ribosomes as they translate a socRNA molecule >100 times allows very precise measurements of ribosome translocation rates on specific mRNA sequences. Second, the unique ability to study either a single or multiple ribosomes on an mRNA enables assessment of ribosome heterogeneity, and provides an opportunity to study ribosome-ribosome interactions as well (Madern et al., 2024). Third, in addition to ribosome speed, socRNAs allow measurement of ribosome processivity, a parameter that has been very difficult to assess with existing assays. Fourth, socRNAs allow uncoupling of translation initiation and elongation, providing opportunities to study translation elongation under conditions of global translation initiation suppression, including stress and viral infection. Fifth, socRNAs can also be used as a sensitive readout for ribosomal frameshifting. Finally, the socRNA method is very easy to implement as a method to study translation elongation because of the high signal intensity of foci and the low temporal resolution required for assessing elongation rates. Thus, we anticipate it can be implemented in cell lines, tissues and potentially even whole organisms, making it a very broadly applicable technology.

### Ribosome processivity

Under our experimental conditions, ribosomes are highly processive, translating on average ∼3 hrs before aborting translation, which corresponds to ∼26,000 codons. Abortive translation can be caused by ribosome frameshifting followed by termination on a stop codon in an alternative reading frame, ribosome recycling, termination on a sense codon or by decay of the socRNA through endonucleolytic cleavage. To understand the effects of a specific sequences on ribosome processivity, it will be important to identify which of the possible mechanisms is causing reduced processivity. Using dual-color socRNAs, frameshifting can be directly assessed. Ribosome recycling by quality control pathways can be assessed through knockdown of quality control proteins (Madern et al., 2024). On the control socRNAs tested here, ribosome frameshifting is the major cause of abortive translation, explaining ∼60% abortive translation. However, for socRNAs containing problematic sequences, including strong pause sequences, other processes may limit ribosome processivity. While a frameshifting rate of 1 in 42,000 codons may seem low, it represents 0.8% of ribosomes on an mRNA of average length in human cells, and >50% of all ribosomes translating the longest human mRNA, Titin. It is important to note that our socRNAs contain non-natural sequences. Native mRNA sequences, especially those of long mRNAs, may have evolved to suppress frameshifting, which will be interesting to investigate in the future.

### eIF4A in translation elongation

By uncoupling translation initiation and elongation, we were able to use socRNAs to identify a novel role for the translation initiation factor eIF4A in translation elongation. eIF4A is thought to promote translation initiation by unfolding RNA structure in the 5’UTR during 43S ribosome scanning. However, eIF4A is expressed at approximately 10-fold higher levels than other translation initiation factors, and recent structural work revealed a second eIF4A molecule at the mRNA entry channel of the small ribosome subunit (Brito Querido et al., 2024), leading to speculation about additional functions of eIF4A. Indeed, recent work showed that eIF4A is also involved in disassembly of stress granules (Tauber et al., 2020). Our work shows that eIF4A is additionally important for translation elongation, as inhibition of eIF4A reduced translation elongation rates, albeit modestly. Somewhat surprisingly, translation elongation rates were not further reduced by introduction of a strong RNA structure in the socRNA. Perhaps even mRNAs that don’t contain obvious hairpin structures are already highly folded (Ruijtenberg et al., 2020), so introduction of a hairpin doesn’t increase the overall thermodynamic stability (ΔG) of the mRNA substantially. Alternatively, eIF4A may have a role in translation elongation independent of its role in unfolding RNA structures, for example in removing proteins from the mRNA (Gentry et al., 2023).

### Heterogeneity in translation elongation rates

In this study we show that individual ribosomes move at distinct speeds during translation. Our results show that distinct translation speeds cannot be explained by technical noise, diverse mRNA sequence, size of the nascent chain or cell-to-cell heterogeneity, leaving three possible explanations; first, it is possible that different socRNAs are differentially modified and that such modifications impact translation speed. However, we feel this is unlikely considering that all socRNAs are transcribed from the same promoter and processed in the same way. Moreover, even if socRNAs are differentially modified, modified nucleotides would need to be decoded extremely slowly to quantitatively explain the observed differences in average translation rate. A second possible explanation is that differences in the sub-cellular compartment of different socRNAs explain the observed translation elongation speed heterogeneity. However, we believe this is also unlikely because all socRNAs analyzed here are tethered to the plasma membrane, making their sub-cellular localization fairly uniform. In addition, socRNA mobility does not correlate with translation elongation speed, suggesting that socRNAs translated at distinct speeds are not present in a confined compartment or anchored to a cellular organelle. Based on these observations, the most likely explanation for our data is that different ribosomes translate the same sequence at distinct speeds due to intrinsic ribosome heterogeneity. Elongation speed heterogeneity might be caused by heterogeneity in rRNA sequence or modifications, which are known to be heterogeneous between ribosomes (Parks et al., 2018), compositional or structural differences in ribosomes, differences in associated proteins (e.g., eIF4A) or damage to ribosomal proteins or RNA. Functional and structural heterogeneity of ribosomes is a field of intense investigation (Gay *et al*., 2022; Genuth and Barna, 2018), and the ability to study translation elongation of individual ribosomes using socRNAs adds a valuable tool to this field.

### Limitations of socRNAs

While the socRNA method has many advantages, it also has a number of potential limitations, both technical and biological, that should be considered carefully. A biological limitation is that socRNAs lack a 5’ cap and poly(A) tail, so any regulation that requires these RNA elements will not be active on socRNAs. The lack of a cap and poly(A) tail can also be leveraged as an advantage under some circumstances, however. For example, lack of these elements prevents canonical RNA decay by XRN1 and the exosome, allowing study of translation in cases where exonucleolytic RNA decay pathways would otherwise have degraded the mRNA. Similarly, socRNAs also lack 3’UTRs, which are known to harbor regulatory elements. While most 3’UTR regulatory elements affect translation initiation and/or mRNA decay, some may affect translation elongation as well. A potential technical concern is that the circular topology of the socRNA may create tension along the RNA, which could affect translation elongation. However, this is very unlikely considering that RNA is an extremely flexible molecule (persistence length ∼1 nm) (Hyeon et al., 2006). Moreover, even linear mRNA may form a circular topology under certain conditions (Vicens et al., 2018). Indeed, we find that translation elongation rates on linear mRNAs and socRNAs are similar. Another technical concern is that the very large nascent chain could potentially slow down ribosome translocation. However, we find no evidence for hindrance of ribosome translocation by the large nascent chains (Figures 5J and 5K). Finally, membrane tethering may position socRNAs in a cellular environment that differs from other parts of the cytoplasm and may affect elongation dynamics. Our previous work examining tethered and untethered mRNAs, however, has revealed that membrane tethering does not affect translation regulation or dynamics (Hoek et al., 2019; Ruijtenberg *et al*., 2020; Yan *et al*., 2016). As a control, membrane tethering can be omitted and socRNAs can be tracked in 3D in the cell, if necessary.

In summary, socRNAs represent a powerful new assay to study translation elongation and will hopefully find widespread use to study the kinetics and mechanisms of regulation of translation elongation.

## Acknowledgements

We thank members of the Tanenbaum for helpful discussions. We also thank Srinaath Narasimhan for help with experiments and Olivia Rissland for critical reading of the manuscript. M.F.M., S.Y., and M.E.T. were supported by the Oncode Institute, which is partly funded by the Dutch Cancer Society (KWF). M.E.T. acknowledges funding from the VIDI (NWO/016.VIDI.189.005). S.Y. acknowledges funding from the European Union’s Horizon 2020 research and innovation program under the Marie Skłodowska-Curie grant agreement no. 101026470. M.B. acknowledges funding from the VIDI (NWO/VI.Vidi.223.169).

## METHODS

### Cell lines

Human U2OS, HEK293T cells used for imaging and lentivirus production were grown in DMEM (4.5 g/L glucose, Gibco) supplemented with 5% fetal bovine serum (Sigma-Aldrich) and 1% penicillin/streptomycin (Gibco). All cells were grown with 5% CO_2_ at 37°C. Cells were confirmed to be mycoplasma negative.

### Plasmids

The sequences of plasmids used in this study can be found in Table S2.

### Cell line generation

To generate cell lines with stable transgene expression, lentiviral transduction was used. Lentivirus was produced in HEK293T cells by transfecting cells at 40 % confluency with a lentiviral plasmid along with the packaging vectors psPax and pMD2 using Polyethylenimine (PEI) (Polysciences Inc). The cell culture medium was replaced 1 day after transfection, and the supernatant containing lentivirus was harvested 3 days after transfection. For lentiviral transduction, U2OS cells were seeded in 6-well plates and virus-containing supernatant was added to cells together with Polybrene (10 mg/mL) (Santa Cruz Biotechnology Inc). Cells were then spin-infected for 100 minutes at 2000 rpm at 37 °C. To generate monoclonal cell lines with homogeneous expression levels of the transgenes of interest, single cells were FACS sorted into 96-well plates.

### Drug treatment

To precisely quantify the number of translating ribosomes on socRNAs, the translation inhibitor puromycin (0.1 mg/mL; ThermoFischer Scientific) was added to cells 1-3 hours after the start of imaging to induce premature nascent chain release. To assess the effect of different translation inhibitors on elongation speed, harringtonine (3 µg/mL; Cayman Chemical), cycloheximide (200 µg/ml), or hippuristanol (5 µM) were added to the imaging medium at indicated time-points (Figures 1 and 4). For studying the kinetics of ribosome-targeting drugs (Figure 3), cycloheximide (25 µg/mL), anisomycin (5 µg/mL), and narciclasine (5 µg/mL) were added to the medium for 15 minutes, followed by subsequent washout through three sequential wash steps during live-cell imaging. MG132 (10 µM) was added in frameshifting assays to prevent any potential decay of (nascent) polypeptides.

### Live-cell microscopy

#### Microscopes

Imaging experiments were performed using a Nikon TI inverted microscope with NIS Element Software equipped with a perfect focus system, a Yokagawa CSU-X1 spinning disc, an iXon Ultra 897 EM-CCD camera (Andor), and a motorized piezo stage (Nanocan SP400, Prior). The microscope was equipped with a temperature-controlled box. A 100x 1.49 NA oil-immersion objective was used for all imaging experiments.

#### Cell culture for imaging

Unless noted otherwise, socRNA imaging was performed by seeding cells stably expressing STAb-GFP, ALFANb-CAAX, and TetR in a 96-well glass-bottom plate (Matriplates, Brooks Life Science Systems) at ∼25% confluency. The next day, the cells were transfected using Fugene (Promega) with a plasmid encoding the socRNAs of interest. Imaging was done the following day by replacing the medium with pre-warmed imaging medium (CO2-independent Leibovitz’s-15 medium (Gibco) containing 5% fetal bovine serum (Sigma-Aldrich) and 1% penicillin/streptomycin (Gibco)). 90 minutes prior to the start of imaging, doxycycline (Dox, 1 µg/mL) was added to the cells to induce socRNA expression. All live-cell imaging experiments were performed at 37 °C.

#### Single-molecule imaging of socRNAs

For live-cell imaging of socRNAs, the x, y positions for imaging were chosen based on the presence of translating socRNAs in cells. Images were acquired every 90-180 sec for 1-4 hours with exposure times for the 488 laser ranging from 50-100 ms. Unless stated otherwise, single z-plane images were acquired with focus on SunTag-GFP foci on the plasma membrane. For experiments in which the GFP fluorescence intensity of individual 24xSunTag arrays was measured, the cells were transfected with a plasmid encoding the 24xSunTag-CAAX protein.

#### Ribosome frameshifting

For imaging of ribosome frameshifting, a monoclonal U2OS cell line stably expressing ALFANb-Halo-CAAX, STAb-GFP, and TetR was used. 1 hour prior to live-cell imaging of frameshifting, cells were incubated with 50 nM HaloJF^549^ for 40 minutes, after which cells were washed twice to remove unbound dye. 15 minutes later, cells were washed once again and positions for imaging were selected. Cells were imaged at 180 sec interval for 4-5 hours using a 488 laser lines (30% LP, 50 ms exposure time) and a 561 laser line (4% LP, 100 ms exposure time).

For the high-throughput assay to determine frameshifting rates of multiple different socRNAs, we induced socRNA expression using doxocycline and added puromycin to cells 4 hours after socRNA induction, leading to the release of all nascent chains. Puromycin was added in this assay to ensure that all foci that were positive for both SunTag and ALFA-tag signal represented bona fide frame-shifting products, rather than two ribosomes translating the same socRNA in different reading frames. MG132 was added together with doxocyline and was present until the moment of imaging to prevent degradation of protein products. HaloJF^549^ was added to cells prior to imaging and later washed out again, as described above. 60-90 minutes after puromycin addition, cells were imaged for 15 minutes at 3 minute interval. Time-lapse imaging was performed to ensure that dual color foci represented single polypeptides labeled in both colors, rather than two polypeptides that co-localized by chance in a single time-point.

### Single-molecule Fluorescence In Situ Hybridization (smFISH)

#### Probe Labeling for Single-Molecule Fluorescence In Situ Hybridization (smFISH)

Single-molecule fluorescence in situ hybridization (smFISH) was conducted following established protocols (Lyubimova et al., 2013; Raj et al., 2008). Forty custom oligonucleotide probes targeting the 5xSunTag socRNA were designed using the Stellaris probe designer available at www.biosearchtech.com (Table S3 for probe sequences). The labeling of the probes was accomplished using ddUTP-coupled Atto633 dyes (AttoTec) in conjunction with terminal deoxynucleotidyl transferase, as previously detailed (Gaspar et al., 2018). Following probe synthesis, purification entailed precipitation of the labeled probes using 100 % ethanol, subsequent washing with 80% ethanol, and final resuspension in nuclease-free water.

#### Probe hybridization

To fix cells for smFISH staining, cells cultured in 96-well glass-bottom plates were first washed once with PBS and then incubated with 4% paraformaldehyde in PBS for 5 minutes at room temperature (RT). Subsequently, cells were subjected to two PBS washes, followed by incubation with 100% ice-cold ethanol at 4 °C for 30 minutes. Cells were then washed twice with a wash buffer (2xSSC and 10% formamide in diethyl pyrocarbonate-treated water at RT). The labeled smFISH probes were diluted to a concentration of 10 nM in hybridization buffer (1% dextran sulfate, 2xSSC, and 10% formamide in diethyl pyrocarbonate-treated water) and added to the fixed cells, followed by probe hybridization within a sealed container at 37 °C for the duration of 16 hours. To wash away unbound probes, cells underwent two washing cycles with wash buffer lasting for 1 hour each at 37 °C. DAPI was included at 1 µg/ml during the second of the two wash cycles. Finally, cells were washed with another 15 min wash step at RT. For imaging, the wash buffer was replaced with imaging buffer (10 mM Tris pH8, 2xSCC, 0.4% glucose, containing both glucose oxidase (Sigma-Aldrich) and catalase (Sigma-Aldrich)). Imaging was carried out at RT.

### Design of socRNAs differing in codon optimality

To design socRNAs encoding the same protein but differing in the codon adaptation index, a custom-written script was used to select codons based on the frequency with which they occur in the human genome. While 11% of the socRNA coding sequence are necessary for RNA circularizaiton, we changed the remaining 89% in the following way:or codon-randomized socRNAs, each synonymous codon was chosen randomly and with equal probability. For each codon in codon-optimized socRNAs, rare synonymous codons were entirely excluded for codon selection, and the remaining codons were randomly selected with equal probability. For each codon in codon-deoptimized socRNAs, a codon out of the 1-2 rarest synonymous codons was randomly selected. Using the approach described above, 5 different socRNAs encoding the same protein were synthesized, with the following CAI scores: 0.67 (codon-randomized #1), 0.66 (codon-randomized #1), 67 (codon-randomized #2), 0.84 (codon-optimized), 0.49 (codon-deoptimized #1), 0.51(codon-deoptimized #2).

The list of rare codons excluded in the codon-optimized socRNA is as follows:

- **Leucine (L):** UUA, UUG, CUU, CUC, CUA
- **Isoleucine (I):** AUA
- **Serine (S):** UCG, AGU
- **Proline (P):** CCC, CCG
- **Threonine (T):** ACG
- **Alanine (A):** GCC, GCG
- **Glutamine (Q):** CAA
- **Arginine (R):** CGU, CGA
- **Glycine (G):** GGU, GGG

The list of codons excluded in the codon-deoptimized socRNA is as follows:

- **Leucine (L):** UUG, CUU, CUC, CUG
- **Isoleucine (I):** AUU, AUC
- **Valine (V):** GTC, GTG
- **Serine (S):** TCT, TCC, TCA, AGT, AGC
- **Proline (P):** CCU, CCC, CCA
- **Threonine (T):** ACU, ACC, ACA
- **Alanine (A):** GCU, GCC, GCA
- **Glutamine (Q):** CAG
- **Arginine (R):** CGC, CGG, AGG
- **Glycine (G):** GGC, GGA, GGG

### Sample preparation for socRNA sequencing

To validate the sequence of socRNAs, RNA was isolated from cells 3 hours after inducing socRNA expression using TRIsure (Bioline). Subsequently, cDNA was synthesized utilizing a gene-specific primer designed to target the 10xSunTag socRNA and Tetro Reverse Transcriptase (Bioline). The resulting cDNA was isolated via column-based purification using the GeneJet Gel Extraction Kit (Thermo Scientific). To generate dsDNA for sequencing using the cDNA as template, three distinct polymerase chain reaction (PCR) reactions were performed to amplify regions which together cover the entire socRNA sequence. Following purification of the PCR products, each PCR product was sent for Sanger sequencing together with the reverse primer used in the corresponding PCR.

## QUANTIFICATION AND STATISTICAL ANALYSIS

### Post-acquisition processing of microscopy data

For all images, flat-field correction was performed using images obtained from concentrated dye solutions (4 µg/mL DyLight™ 488 NHS Ester for 488 laser line, and 40 µg/mL Alexa Fluor™ 555 NHS Ester for 561 laser line) and dark current images.

For experiments investigating single ribosome heterogeneity in translation elongation rates (Figure 5), we wanted to correct for possible drift in the z-direction, since foci intensity changes slightly even when foci move <100 nm in z. Therefore, 9 z-slices were acquired with a 2 µm total z distance surrounding the GFP foci. Foci intensity was measured in each z slice to acquire a Gaussian profile of GFP foci in the z-direction. To capture the maximum intensity of individual GFP foci at each time point, we first summed the intensity values of 3 adjacent slices across the different z-positions at each pixel, resulting in total 7 summed intensity values at each pixel (This approach is conceptually similar to a moving average over a sliding window length of 3). Then, we used the maximum value among the 7 summed-values for each pixel to generate a maximum intensity projection image at each time point. The reason we used the maximum value of the summed values of 3 adjacent slices instead of a maximum intensity projection is to avoid maximizing the background intensity from the non-GFP foci area.

### Tracking and intensity measurements of socRNA foci

For tracking and fluorescence intensity measurements of socRNAs, we used the ‘TransTrack’ software package as previously described) (Boersma *et al*., 2019). All resulting traces underwent manual curation to ensure accuracy.

To correct for photobleaching of membrane-tethered GFP-foci, we used GFP intensity time traces from foci exhibiting no increase in intensity over time, referred to as ‘non-translating traces’ (Figure S4C), which were acquired in the same imaging experiments. The decrease in fluorescence intensity of these non-translating GFP foci over time was fit with a single exponential decay function to determine the bleaching rate. All GFP foci intensities were then corrected for the photobleaching (Figure S4D). We used the photobleaching corrected non-translation traces as ‘plateau traces’ for further analysis. We found that bleach correction on GFP foci rather than whole cell fluorescence is essential, as GFP foci bleach faster than the whole cell, because only a small region of the cell in the z-direction is excited by laser light, while GFP foci stay within the excitation focus plane throughout the experiment and thus bleach faster than the whole cell fluorescence.

### Quantification of smFISH results

To assess the co-localization of smFISH spots with socRNA translation sites (Figure S1A), socRNA translation sites were first imaged and tracked over time, followed by smFISH staining of the same cells. Combining live-cell imaging and smFISH of the same cells allowed us to determine co-localization of smFISH RNA foci for both translated and non-translated socRNAs. After live-cell imaging, cells were quickly fixed, smFISH staining was performed and the same cells were imaged again to co-localize smFISH signal and socRNA translation signal, which was preserved after fixation. Intensity threshold-based masks were generated for each SunTag-positive object, and the presence or absence of a co-localizing smFISH spots was scored using Fiji. To control for chance co-localization, the far-red channel (smFISH signal) was rotated by 90 degrees relative to the green channel (socRNA translation products), and the same analysis was carried out again.

### Translation elongation rates of single ribosomes on socRNAs

To determine the number of translating ribosomes per socRNA, puromycin (0.1 mg/mL) was added to cells at the end of the imaging experiment and the number of ribosomes was determined by counting the number of splitting GFP foci after puromycin addition (Figures 1J-1L). The elongation rate of ribosomes on socRNAs was determined by fitting a linear function to GFP intensity time traces to extract the slope of intensity increase phase before puromycin addition. The slope was then divided by the number of ribosomes to determine the translation elongation speed per ribosome. To convert rates of GFP intensity increase into the unit of amino acids translated per second, we first determined the intensity of a single GFP molecule under our experimental settings. To achieve this, we measured the intensity of individual ‘mature’ SunTag proteins containing 24 repeats of the SunTag peptide fused to a CAAX motif (24xSunTag-CAAX) using the same settings as those used in the imaging experiment (Figures S6F and S6G). We divided the average intensity of 24xSunTag-CAAX foci by 24 to obtain the intensity of a single GFP molecule. Using the intensity of a single GFP molecule, we could calculate the number of SunTag epitopes synthesized per unit of time for translating socRNAs. Next, for each socRNA, we calculated the average number of codons that need to be translated for the synthesis of one SunTag epitope; we determined the number of codons for the translation of a full cycle for each socRNA, and the number of SunTag epitopes synthesized upon translation of the socRNA once (equal to the number of SunTags encoded in a socRNA, 5 or 10, unless noted otherwise). Based on the number of codons in one full cycle of socRNA translation and the number of SunTag epitopes encoded in a socRNAs, we calculated the elongation rate in amino acids per second.

### Calculating ribosome pause time

To determine ribosome pause time on socRNAs encoding a pause sequence, we determined the average elongation rates (i.e., the total time to complete translation of one full circle, which represents the time needed to translate the non-pause sequence plus the pause time on the pause sequence) of single ribosomes as described in the paragraph above. We then subtracted the average translation time for one cycle of translation of a matched socRNA lacking the pause sequence to obtain the pause duration per cycle.

### Calculating off-rates of translation elongation inhibitors

To quantitatively assess binding kinetics of elongation inhibitors, we generated intensity time traces of translated socRNAs from cells treated with elongation inhibitors, followed by inhibitor washout. To identify the moment of unbinding of the translation inhibitor, we wished to identify the precise moment in time when the GFP intensity time trace transitions from a plateau to a positive slope. To identify this transition point, a custom-written python script was applied, which employed two distinct linear regression models to fit the intensity time trace. The least squares method was used to find an optimal fit. The linear regression model for the first half of the intensity time trace was constrained to have a slope of zero (representing the time when the inhibitor is still bound to the ribosome), while the linear regression model for the second part of the intensity time trace needed to exhibit a positive slope (representing the time when the inhibitor was released from the ribosome and translation had resumed).

### Quantification of ribosome processivity

To determine the number of codons translated by individual ribosomes on socRNAs, we tracked GFP intensity time traces for translated socRNAs and determined the moment when the GFP intensity stopped increasing for individual translated socRNAs. For socRNAs translated until puromycin addition, we noted the last frame before puromycin addition as the last time-point in which translation was detected. We then measured for each individual socRNA the GFP foci intensity at the last time-point of translation and calculated the total number of codons translated during the experiment based on this final time-point GFP intensity, as described in *Translation elongation rates of single ribosomes on socRNAs*. The fraction of translated socRNAs remaining was then plotted against the total number of codons translated in Kaplan-Meier survival plots.

### Contribution of cell-to-cell heterogeneity to single ribosome elongation rate heterogeneity

To quantify the contribution of cell-to-cell heterogeneity to single ribosome elongation rate heterogeneity, we randomly selected two ribosomes translating two different socRNAs within the same cell (Figure 5E) and employed an approach used for the noise decomposition into intrinsic and extrinsic components, which have orthogonal contribution to total noise (Figure 5F) (Elowitz et al., 2002; Swain et al., 2002). In brief, the total noise (defined as the standard deviation divided by the mean) in the ribosome elongation rates (Figure 5C) can be separated into two components: intrinsic noise (e.g., variation between ribosomes) and extrinsic noise (e.g., variation between cells). Extrinsic noise corresponds to the data spread parallel to the diagonal line on the scatter plot showing the speed of the two randomly selected ribosomes from the same cell (Figure 5F). On the other hand, intrinsic noise is represented by the data spread perpendicular to the diagonal line on the scatter plot. Intrinsic, extrinsic, and total noise were defined as follows:

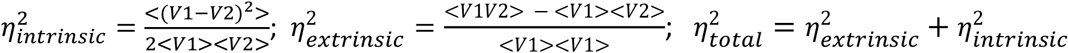

where *V1* and *V2* represent the elongation rates of either of two ribosomes randomly picked from the same cell, respectively. Angled brackets denote means over the cell population. Based on this approach, we calculated that only ∼22% of the ribosome speed variation originates from extrinsic noise, i.e., cell-to-cell heterogeneity, with the majority of variation originating from intrinsic noise.

### Identifying transient pauses in GFP intensity time traces from single ribosomes translating socRNAs

To identify pauses within single ribosome intensity time traces (Figures 5L-5N), the raw intensity traces (black line in Figure 5L) were first smoothed using a moving median to eliminate outlier data points. Subsequently, a moving average was applied to further smooth the data (red line in Figure 5L). Following this, the first derivative, which represents the differences between adjacent intensities, was calculated (black line in Figure 5M). Pause identification was performed using a hidden Markov model (vbFRET algorithm (Bronson et al., 2009)) with a maximum of 2 states, with default settings of the algorithm for other parameters. The threshold for the value of pause state (i.e., the derivatives between adjacent intensities) was set at a value lower than (mean-2*standard deviation) of the histogram of the fitted states in the negative control to ensure minimal false positive calling of pauses. The negative control traces representing the experimental noise (Figure 5B) were generated from the ‘plateau traces’. As a positive control, we simulated intensity time traces with pauses of known duration. For this, we used plateau traces which we transformed using a constant value to mimic increasing traces (i.e., translated socRNAs). We then added a 10-time-point (15 min) pause (i.e., no increasing intensity for 15 min) in the middle of the increasing intensity traces.

### High throughput assay for determining ribosome frame-shifting rates

To determine the frame-shifting rates for various different socRNAs in a high through-put manner, co-localization of ALFA-tag and SunTag spots was assessed in cells. The Fiji plugin ‘ComDet’ was used to determine co-localization of spots from both fluorescence channels. frame-shifting products were called if ALFA-tag and SunTag foci co-localized for 15 consecutive minutes) of live-cell imaging (3-minute interval). In addition, fluorescent intensities of all spots in the ALFA-tag channel (which represents the main frame) were measured to subsequently calculate how many codons each ribosome had translated before frameshifting occurred. Using the 24xSunTag-CAAX reporter described above and a socRNA encoding an equal number of ALFA-tag and SunTag epitopes, we could normalize fluorescent intensities of foci in the two channels to the absolute amount of fluorescent proteins, and thus the number of SunTag/ALFA-tag epitopes, that have been translated. Using the ALFA-tag intensity information from both frameshifted and non-frameshifted proteins, we constructed survival plots correlating the number of translated codons to the fraction of ribosomes that have undergone frameshifting. To calculate the average frameshifting rate per codon, a single-exponential decay function was fit to our survival curve.

### Mobility of translating socRNAs

To acquire the x, y coordinates of individual translating socRNAs at each time, we tracked the socRNAs using TransTrack (Boersma *et al*., 2019) with 90 sec time intervals. Using the x,y position information of foci at each time point, we calculated the mean squared displacement as a measure of the mobility of translating socRNAs.

### Statistical analyses and generation of graphs

All graphs were generated using Prism GraphPad (v9) or in python 3.10 using Matplotlib. Details of statistical tests for each graph are explained in figure legends.

## THEORETICAL MODELING OF RIBOSOME ELONGATION RATES HETEROGENEITY

### Main results

We consider the single ribosome translation traces, and investigate the heterogeneity observed in the estimated translation elongation rates 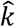. In particular, we ask the question if the heterogeneity in 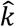 can be explained by a combination of technical noise and noise from the stochastic movement of the ribosome, or whether the translation elongation rates themselves must differ among ribosomes to explain the data. In this section we outline the main results, and in next section we provide the technical details.

To characterize the technical noise, we use traces that were not increasing in GFP intensity (“plateau traces”). These traces do not contain noise caused by stochastic movement of the ribosome. We find that the technical noise is well-described by Gaussian white noise, with a variance that scales linearly with the mean spot intensity. Further, we model the ribosome movement along the socRNA as a homogenous one-dimensional Poisson process with a mean rate *k*. From our description of the system, we can estimate both analytically and through simulation the expected heterogeneity in estimated translation elongation rates 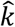, and compare it to the experimental data (Figures 5D and S3A).

Figure S3A shows a scatterplot of the estimated starting length of the polypeptide chain 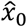 and the estimated translation elongation rate 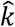. These estimators are obtained by performing a least-squares linear regression on the single-ribosome translation traces (“moving traces”). Appropriately adjusting for units, 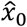 and 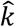 correspond to the *y*-intercept and slope of the linear fit, respectively. On the same plot, we superimpose the distribution 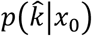, which we obtained analytically. This shows the distribution of ⟩ that we expect from our model given a particular starting length *x*_0_.

In Figure 5D, we show the histogram of translation elongation rates 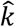 estimated from the moving traces in red. To compare it to our analytical prediction, we integrate out dependence on 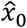 by computing the distribution 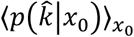. This distribution is shown as the blue line in Figure 5D. As an internal consistency check (and to verify our analytical results) we simulated the ribosome movement and added technical noise. The blue histogram in Figure 5D shows the spread in translation elongation rates 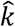 from the simulation, and indeed the analytical distribution matches our simulation results. We observe that the spread in estimated translation elongation rates 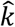 is significantly wider in the data than would be expected from technical noise and noise from stochastic movement alone. This indicates that there are intrinsic differences between the mean translation elongation rates *k*of different ribosomes.

On a final note, we should consider the possibility that modelling the ribosome movement as a Poisson process is invalid. In particular, the ribosome is known to cycle through a series of internal protein configurations between successive steps along the RNA, which can cause the number of steps in a given time interval to no longer be Poisson-distributed. A more detailed description of ribosome kinetics from existing models could be incorporated. However, we are in a regime where the central limit theorem suppresses noise caused by the stochastic movement of the ribosome, and the noise is dominated by technical noise. Hence, we do not expect that choosing different movement statistics for the ribosome will significantly impact the conclusions drawn here.

### Theoretical methods overview

Here we provide the technical details for the result shown in the above section. Our goal is to find out whether the distribution of estimated translation elongation rates 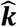 can be explained by a combination of technical noise and noise from the stochastic movement of the ribosome, or whether the rates themselves are heterogenous. We begin by characterising the noise in the experiment in *Characterizing the noise in the system*, which we use to build a stochastic description of the system. Finally, in *Distribution of ribosome translation rates*, we use least-squares regression to obtain the ribosome translation elongation rates from the moving traces and compare the results we get to those expected analytically from our stochastic picture.

### Characterizing the noise in the system

We consider two sources of noise. Firstly, we consider noise due to the stochastic movement of the ribosome. We choose to model the movement as a one-dimensional Poisson process, but we motivate that our results are not model-specific. Secondly, we fully characterize the technical noise using the plateau traces that contain no noise from the movement of the ribosome. Finally, we use our results to formulate a stochastic description of the system.

#### Ribosome movement

To determine how the stochastic movement of the ribosome contributes to the noise, we require a model for its movement statistics. A kinetic description of ribosome translation can be complicated, involving transitions through multiple configurations of the ribosome between each step and recruitment of the appropriate proteins (Rudorf, 2019; Rudorf and Lipowsky, 2015). However, in the experiment, spot intensity is sampled every 1.5 minutes. During this time ∼200 codons have been traversed by the ribosome on average. Hence, by the central limit theorem, the statistics of the ribosome movement becomes Gaussian, and we will see that finer details of the kinetic description are averaged out in this regime.

We begin by assuming that the ribosome movement *x*(*t*) can be modelled as a homogenous Poisson process with a mean rate *k* and initial condition *x*(0) = *x*_0_ (Figure S3B). We can write:

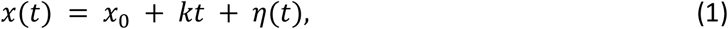

where *η*(*t*) is the noise, which is Gaussian due to the central limit theorem. We have by construction:

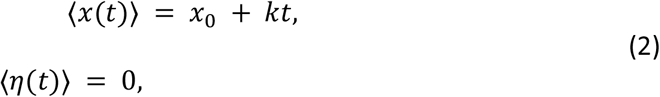

where the angled brackets ⟨⟩ denote the ensemble average. Due to the Poisson statistics, the variance of this process will scale as its mean, meaning that

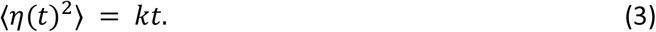

We should consider the possibility that the movement of the ribosome may occur at a mean rate *k* but is *not* a Poisson process. One can, in principle, introduce a more detailed kinetic description for ribosome translation and investigate how this impacts the noise. For example, the ribosome is known to transition through multiple internal protein configurations between each step (Behrmann *et al*., 2015; Rudorf and Lipowsky, 2015). Under the constraint that the ribosome moves with an overall rate *k*, one can show that introducing additional (Poisson-distributed) intermediate transitions between each ribosome step will decrease the variance compared to equation 3. One could also consider transitions to a “pausing” state due to, for example, kinetic proofreading. Such processes could increase the variance in equation 3 while keeping the mean rate *k*fixed. To proceed, we use a more general argument to argue that due to the central limit theorem, we can continue our analysis without subscribing to a specific kinetic description.

Let the time taken for a ribosome to take one step be denoted by a random variable *T*^(1)^. Then, we can define the mean time for one step to occur with E[*T*^(1)^] = 1/*k*, and its variance with 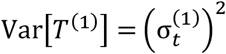. Next, we assume the ribosome steps are independent. Using the central limit theorem, one finds that the noise *η*(*t*) in the movement of the ribosome is Gaussian, with a variance that is given by:

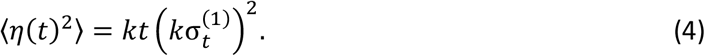

One quick way to see this is by propagating the fluctuations in the time between steps 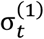 to fluctuations in the ribosome position 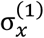, using 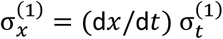. After *kt* steps, we get 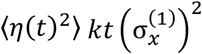 for the variance and equation 4 follows. One can check that setting 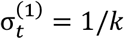 recovers the Poisson case above.

We notice that due to the central limit theorem, the only features from the distribution for *T*^(1)^ that emerge are the mean rate *k* and variance 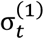. Hence, as we see from equation 4, choosing a particular kinetic description only enters our analysis by scaling the noise *η*(*t*). In section *Stochastic description of the system*, we will see that the technical noise *ξ*(*t*) dominates *η*(*t*). Given that *η*(*t*) is sub-dominant, we do not expect that choosing different movement statistics will impact the conclusion that the ribosome movement occurs at heterogeneous rates.

#### Technical noise

To characterise the technical noise, we use the “plateau traces”. Here, we have no noise from stochastic movement of the ribosome, and hence the intensity *I*(*t*) should fluctuate around a constant value, which we can denote by *ax*_0_:

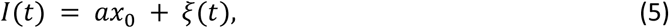

where *a* = 1/64.2 a.u. per amino acid, *ξ*(*t*) is the technical noise, and *x*_0_ denotes the (here unchanging) length of the polypeptide chain. We have:

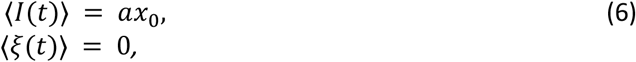

and hence

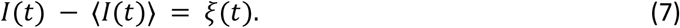

We can therefore gain direct insight into the technical noise by considering deviations from the mean intensity, as in equation 7.

The data is discretely sampled from the continuous system in equation 5. Each plateau trace *I*_*j*_ is sampled at *N* discrete times *t*_*i*_, with *i* = 1, …, *N*. We denote the estimated mean intensity from the *j*th plateau trace as *Î*_*j*,0_ = ∑_*i*_ *I*_*j*_(*t*_*i*_)/*N*. From the data, we notice that the noise is Gaussian. Figure S3C shows a histogram in blue of the residuals, *r*_*j*_(*t*_*i*_) = *I*_*j*_(*t*_*i*_) − *Î*_*j*,0_, from all plateau traces, normalised to unit variance. A normal distribution with unit variance, shown in red, provides a very good fit. To see if the technical noise has correlations, we compute the autocorrelation for the residuals from each plateau trace. Figure S3D shows a superposition of all the autocorrelation functions, which displays a sharp, central peak. Hence the noise is approximately white. Finally, we have to consider how the variance scales with the spot intensity *Î*_0_. We can see in figure S3E that the variance scales linearly with the spot intensity. Combining these observations, the correlation function of the technical noise is given by:

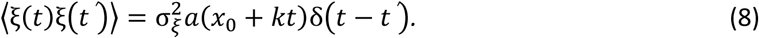

Where 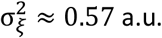 a.u., corresponding to the slope of the line in figure S3E. We have also implicitly used the fact that fluctuations in *x*(*t*) are small compared to ⟨*x*(*t*)⟩.

#### Stochastic description of the system

In order to model the SunTag intensity observed in the experiment, we need to combine our model of the ribosome movement with the technical noise. As above, we denote the observed spot intensity by *I*(*t*), measured in units of GFP fluorescence intensity (a.u.). The number of codons traversed by the ribosome is given by *x*(*t*), and the additive technical noise is denoted by *ξ*(*t*). The observed intensity can then be written as:

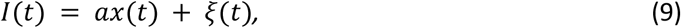

where *a* = 1/64.2 a.u. per amino acid. If the starting position of the ribosome (or, equivalently, the starting length of the polypeptide chain) is *x*(0) = *x*_0_, we can write

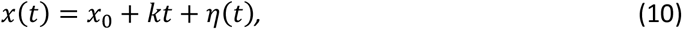

where *η*(*t*) captures Gaussian noise from the stochastic movement of the ribosome, as described in section *Ribosome movement*.

Substituting *x*(*t*) from equation 10 into equation 9, we can describe the system in continuous time with:

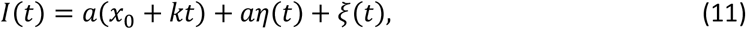

where the covariances of *η*(*t*) and *ξ*(*t*) are given by

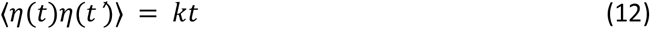

for *t* ≤ *t*′, and

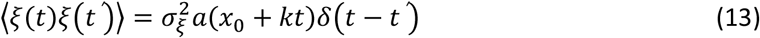

as we saw in sections *Ribosome movement* and *Technical noise*. Further, we assume that the noise terms are independent, i.e. ⟨*ξ*(*t*)*η*(*t*′)⟩ = 0.

Next, we compare the size of fluctuations in *I*(*t*) due to the ribosome movement to those due to the technical noise. From equation 11, one can show that the typical size of the fluctuations in the spot intensity *I*(*t*) is given by:

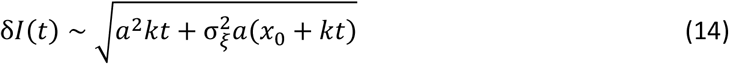

The first term under the square root in equation 14 is suppressed by an extra factor of *a*. Hence, in our model, the fluctuations are dominated by technical noise. This is also true for the data, which one can verified by looking at the residuals in the single-ribosome translation traces.

### Distribution of ribosome translation rates

In this section, we consider the single-ribosome translation traces. Firstly, in section *Obtaining translation rates from data*, we explain how we estimate the translation rates from the data. Next, in section *Obtaining translation rates from the stochastic model*, we show how to obtain the distribution of ribosome translation rates that one would expect analytically, given our stochastic description of the system. The goal is to compare the heterogeneity in the translation rates from the data to the analytical prediction.

#### Obtaining translation rates from data

Given a particular intensity trace, we would like to estimate *x*_0_ and *k*. We denote their respective estimators as 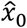 and 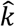. To do so, we perform a least-squares regression to fit a straight line through each trace. Appropriately adjusting for units, the slopes of these lines correspond to an estimate of the translation rate 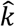, and the ⟩ -intercept corresponds to an estimate of the starting length 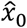. A scatterplot of 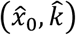 is shown in Figure S3A. Finally, the histogram showing just the distribution of estimated translation elongation rates 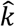 is shown as the red histogram in Figure 5D. The details of the least-squares estimator used to perform the regression are shown in the next section.

#### Obtaining translation rates from the stochastic model

Each trace is sampled at *N* discrete times *t*_*i*_, where *i* = 1, …, *N*. The discrete counterpart of equation 11 can then be written:

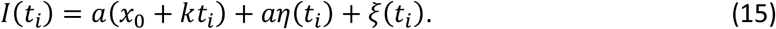

Here, *I*(*t*_*i*_) contains the measured spot intensity at time *t*_*i*_. The terms *η*(*t*_*i*_) and *ξ*(*t*_*i*_) contain noise from the stochastic movement of the ribosome and technical noise, respectively.

To estimate 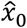 and 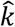, we use a least-squares regression. To help us, we define the *N*-by-2 matrix:

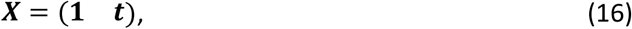

where **1** is an *N*-dimensional vector of ones and ***t*** is a vector with elements *t*_*i*_. Further, we define the parameter vector:

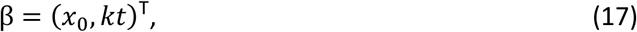

such that equation 15 can be written in vector notation as:

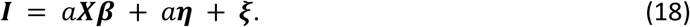

Next, we note that the covariance matrices are given by:

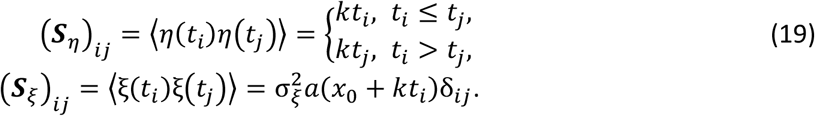

This follows directly from the continuous equivalent in equations 12 and 13. Given that **ξ** and ***n*** are independent and Gaussian, we have

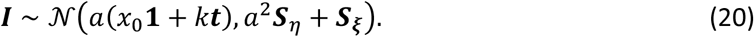

For a given trace **I**, can estimate 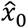 and 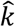 in the least-squares sense by optimising the quantity ℒ(β) = ∥***I*** − *a****Xβ***∥^2^. Setting *∂*ℒ/*∂*β = 0, we obtain the estimator:

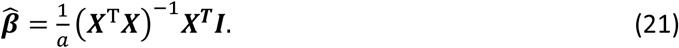

⟩We applied the least-squares estimator in equation 21 to each of the single-ribosome translation traces to obtain the scatterplot in Figure S3A, as outlined in the previous section.

Next, we want to compare what the distribution of estimators 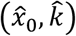 would look like for our stochastic model. To do so, we have to examine the distribution of the estimator 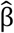 itself. This follows straightforwardly from using equations 20 and 21:

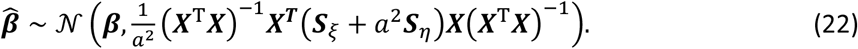

We superimpose the result from equation 22 in Figure S3A (red). Specifically, on Figure S3A we superimpose the distribution 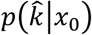. This shows, for a given *x*_0_, the expected distribution of translation rates 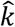. On top of Figure 5D, we plot the distribution of estimated rates 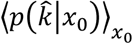 We can clearly see in Figure 5D that technical noise paired with noise from a homogeneous Poisson process is not sufficient to explain the spread in translation rates observed. This indicates that ribosomes are likely to move at different rates.

To test our analytical results, and as an internal consistency check, we simulated the ribosome movement according to our model in equation 11. The simulated traces were generated to have the same length and starting intensities as the real traces, for fair comparison. The blue histogram in Figure 5D shows the distribution of translation elongation rates obtained from the simulated traces. We can see that the analytical result (blue curve) describes the histogram well, and our analysis is therefore internally consistent.

---

**Figure S1.**
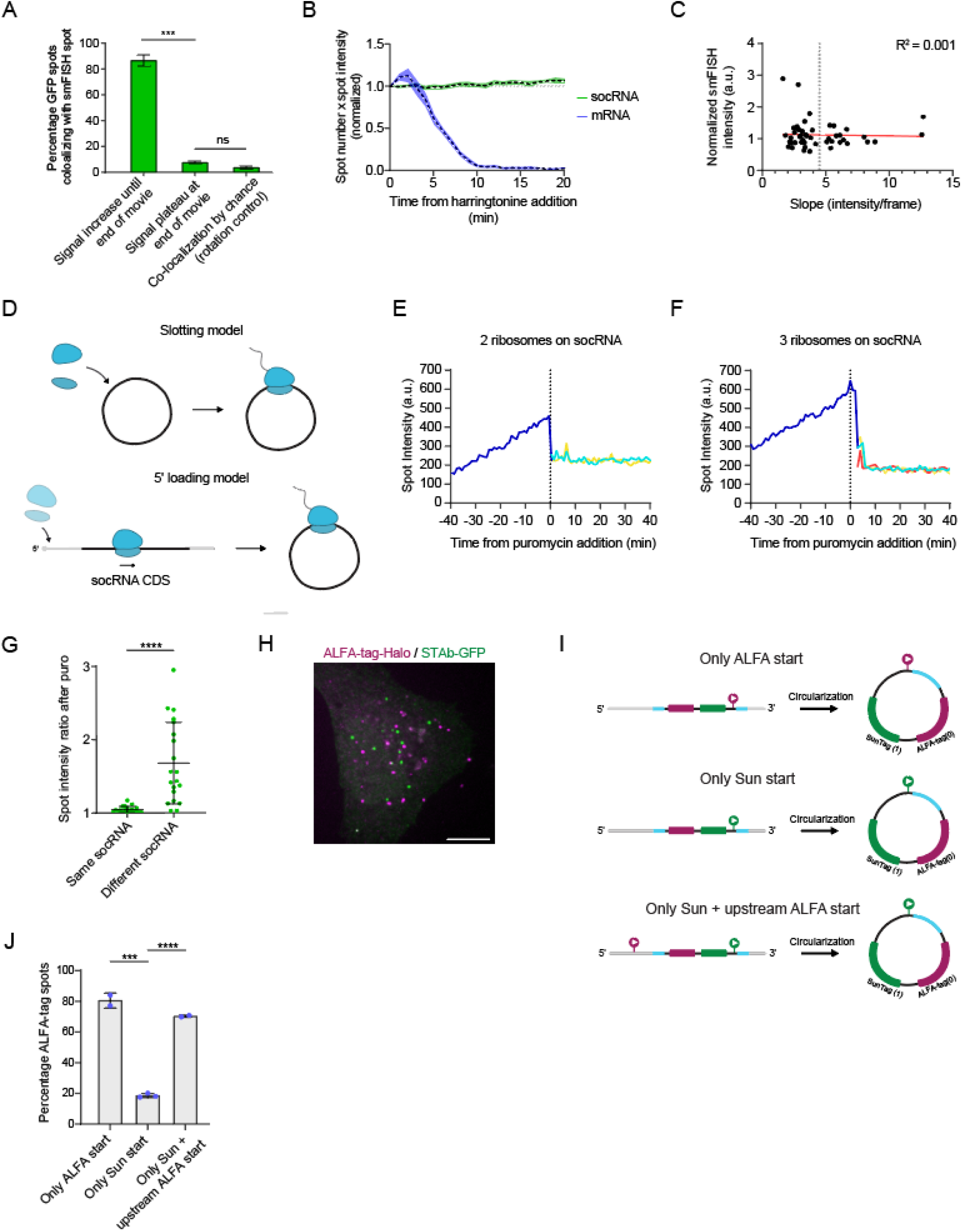
Supplement to Figure 1. Controls for the socRNA translation imaging approach. A) Cells expressing STAb-GFP and the socRNA were followed by time-lapse analysis and GFP intensity of foci was measured over time. After live imaging, cells were fixed and socRNAs were stained by smFISH. Co-localization of GFP translation foci and smFISH foci was assessed for GFP foci that were increasing in intensity at the end of the time-lapse movie (left bar), or for foci that were not increasing in intensity (middle bar). As a control for random co-localization, the image of one channel was rotated and co-localization was assessed (right bar). B) U2OS cells stably expressing STAb-GFP were transfected with either a socRNA (green line) or a linear encoding 24 copies of the SunTag (blue line) and imaged by time-lapse microscopy. Cells were treated with harringtonine and the intensity of translation site foci was measured over time. Dashed lines represent mean values and shaded regions standard error of the mean. C) Cells were treated as in (A) and the smFISH foci intensity was plotted against the GFP intensity increase slope. Note that the smFISH intensity was similar for translating socRNAs that show a much higher slope, indicating that the increased slope is not caused by coincidental co-localization of two or more socRNAs translated by single ribosomes. Dashed gray line separates socRNAs translated by single ribosomes (left of line) from socRNAs translated by multiple ribosomes (right of line) as determined in Figure 1M. D)Schematic depicting two possible models by which ribosomes could be loaded onto socRNAs. In the first model, the “Slotting model”, ribosomes are directly slotted onto socRNA. In the second model, the “5’ loading model”, ribosomes are first recruited to the 5’end of the linear precursor RNA in a cap-dependent mechanism. While the ribosome is translating the coding sequence of the linear pre-cursor RNA, the internal section of the linear RNA becomes circularized, trapping the ribosome in the socRNA. E-F) Cells expressing STAb-GFP and a socRNA were followed by time-lapse analysis and GFP intensity of foci was measured over time. Cells were treated with puromycin at t = 0 to release all the nascent chains from the socRNA. Representative intensity time trace of a socRNA translated by two ribosomes (E) or three ribosomes (F). After puromycin addition two or three new foci are formed (colored lines) that have identical intensities, indicating that all ribosomes translating the same socRNA initiated translation at the same time. G) Relative intensity differences of spots originating from the same socRNA after puromycin treatment. As a control, we compared intensities of spots originating from different socRNAs. Only translated socRNAs that split into 2-3 foci upon puromycin treatment were included in the analysis. H) Representative image of cell line expressing STAb-GFP and ALFANb-Halo transfected with socRNA shown in (I). I) Schematic of reporters used in (J). SunTag and ALFA-tag are encoded in distinct reading frames. Colored arrowheads indicate the frame (magenta = ALFA-tag, green = SunTag) in which the AUG start site is encoded. Cyan regions represent ribozyme sequence. The RNA region in between the two ribozymes will end up in socRNA after RNA circularization. In the top and middle socRNA, the AUG is positioned within the socRNA, while the bottom reporter, there is an additional AUG positioned in the 5’UTR of the linear reporter, which is not included in the socRNA after circularization. J) The three socRNAs shown in (I) were transfected into cells expressing STAb-GFP and ALFANb-Halo. The number of SunTag and ALFA-tag foci was scored, and the percentage of ALFA-tag foci is plotted. Scale bars, 10 μm (H). All error bars represent standard deviations. ***, **** denotes p-values < 0.001, 0.0001, respectively (t-test).

**Figure S2.**
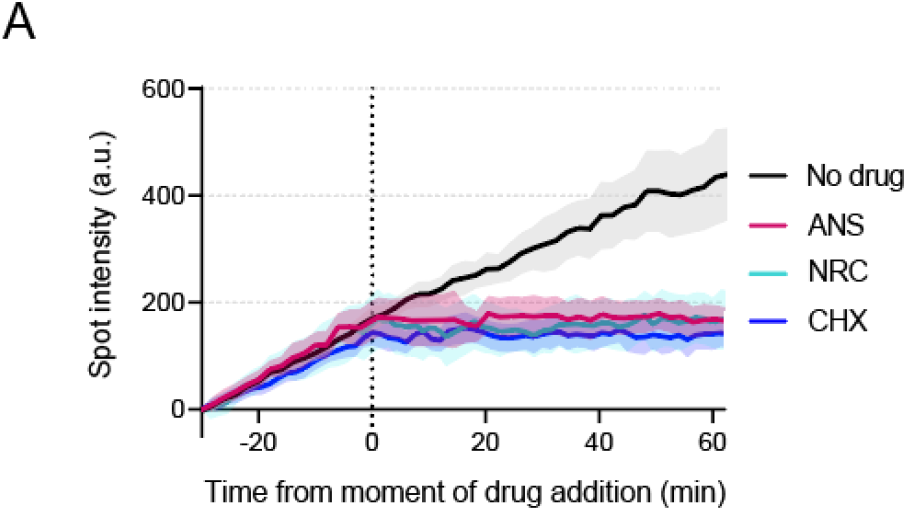
related to Figure 3. Controls for investigating elongation inhibitor kinetics. A) U2OS cells were treated with the translation elongation inhibitors explored in Figure 3. Without drug washout, translation elongation does not resume after drug addition.

**Figure S3.**
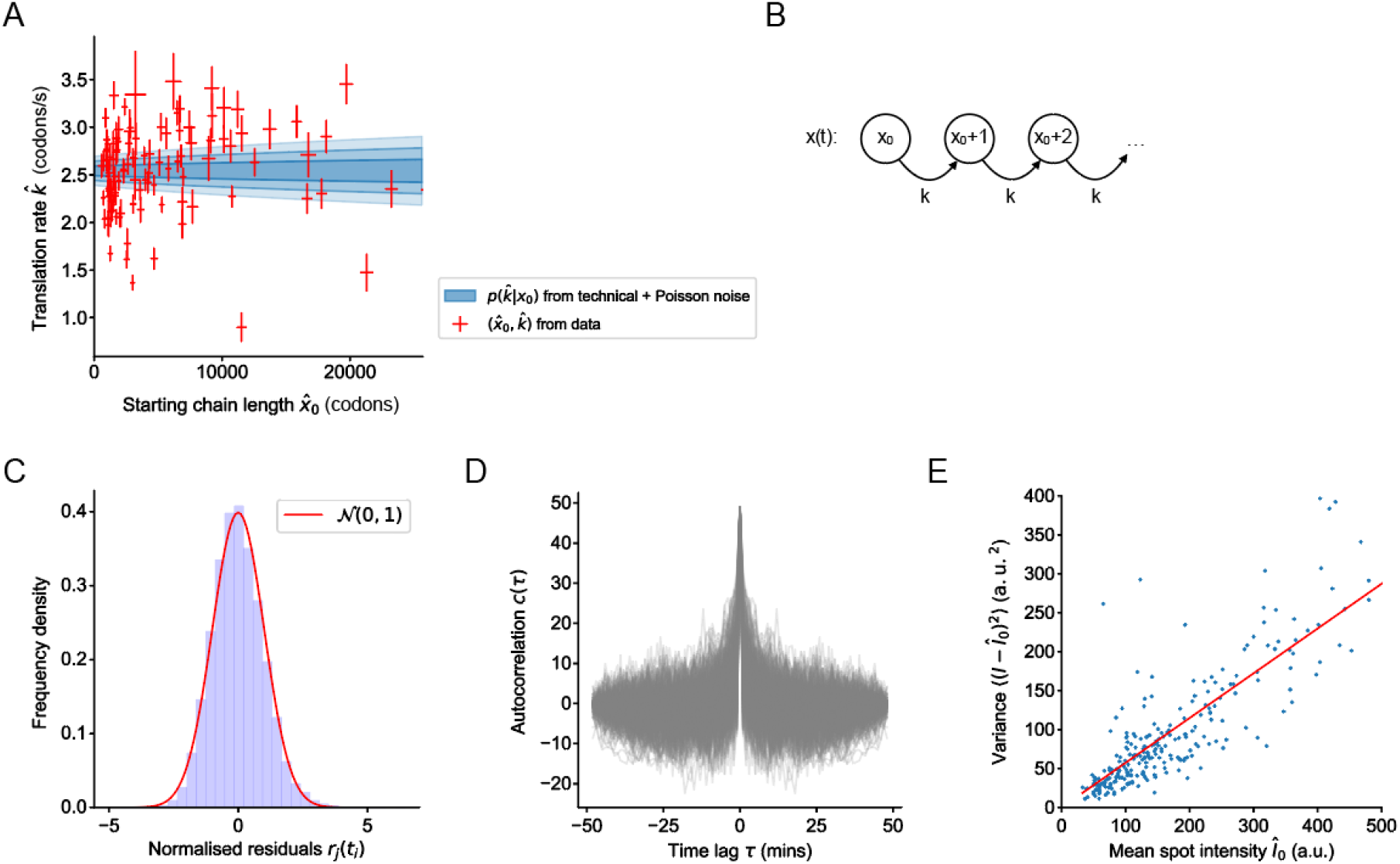
related to Figure 5. Theoretical models for single ribosome elongation rate heterogeneity. A) A scatterplot of estimators 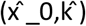, estimated from the single ribosome translation traces. The analytical prediction 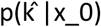 from the model is shown in blue. The darkest shade of blue corresponds to σ, the next lighter shade to 2σ, and so forth. B) A Poisson counting process x(t) with mean rate k amino acids per second and initial condition x(0)=x_0. The number of steps taken by the ribosome in a given time interval is assumed to be Poisson-distributed. C-E) The technical noise can be described by Gaussian white noise. C) A histogram of residuals r_j (t_i)=I_j (t_i)-I^^^_(j,0), normalised to unit variance, from all plateau traces shows that the technical noise is Gaussian. D) The autocorrelation functions c_j (τ)=∑_i▒〖r_j (t_i+τ) r_j (t_i) 〗 are sharply peaked at τ = 0, implying that there are no temporal correlations; this means the noise is white. E) The variance of the technical noise scales linearly with the mean spot intensity.

**Figure S4.**
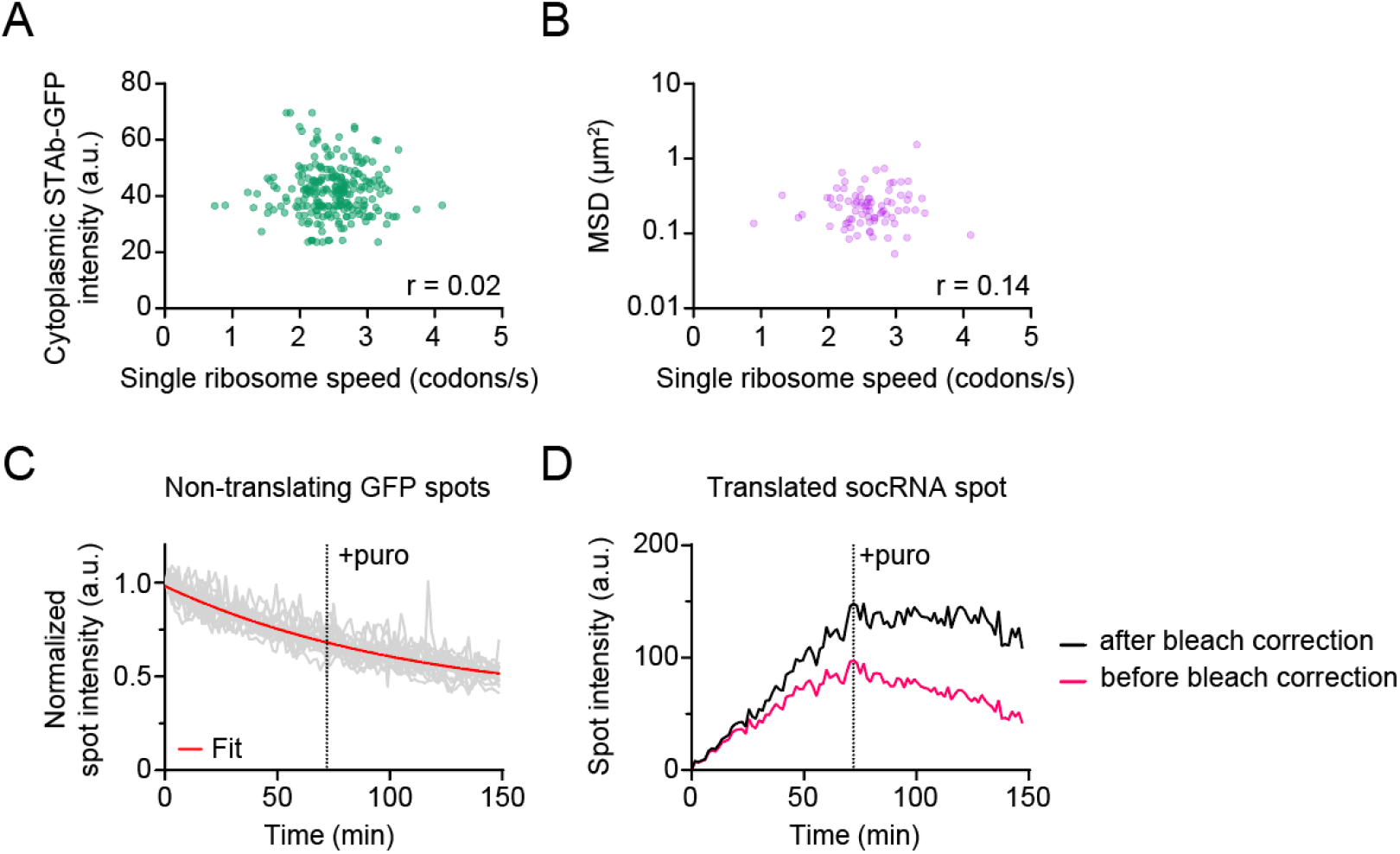
related to Figure 5. Correlation between single ribosome elongation rates and different parameters. A) U2OS cells stably expressing STAb-GFP were transfected with indicated socRNAs and imaged by time-lapse microscopy. Single ribosome translation speeds were calculated and plotted against the expression levels of the STAb-GFP in single cells. No correlation between STAb-GFP expression levels and translation elongation rates was observed. B) Cells were imaged as in (A) and the mobility (mean squared displacement, MSD) of the same translated socRNAs was assessed. Each dot represents one socRNA. No correlation between socRNA mobility and translation elongation rates was observed. C) To correct for Photobleaching, GFP intensity time traces of non-translating GFP foci was measured. Red line represents single exponential decay fitting result that was used to correct for photobleaching for all GFP intensity time traces. D) Example of photobleaching correction for intensity time trace of translated socRNAs. We corrected photobleaching using the value acquired in (C). Note that after bleach correction GFP intensity showed a plateau upon puromycin treatment, as expected.

**Figure S5.**
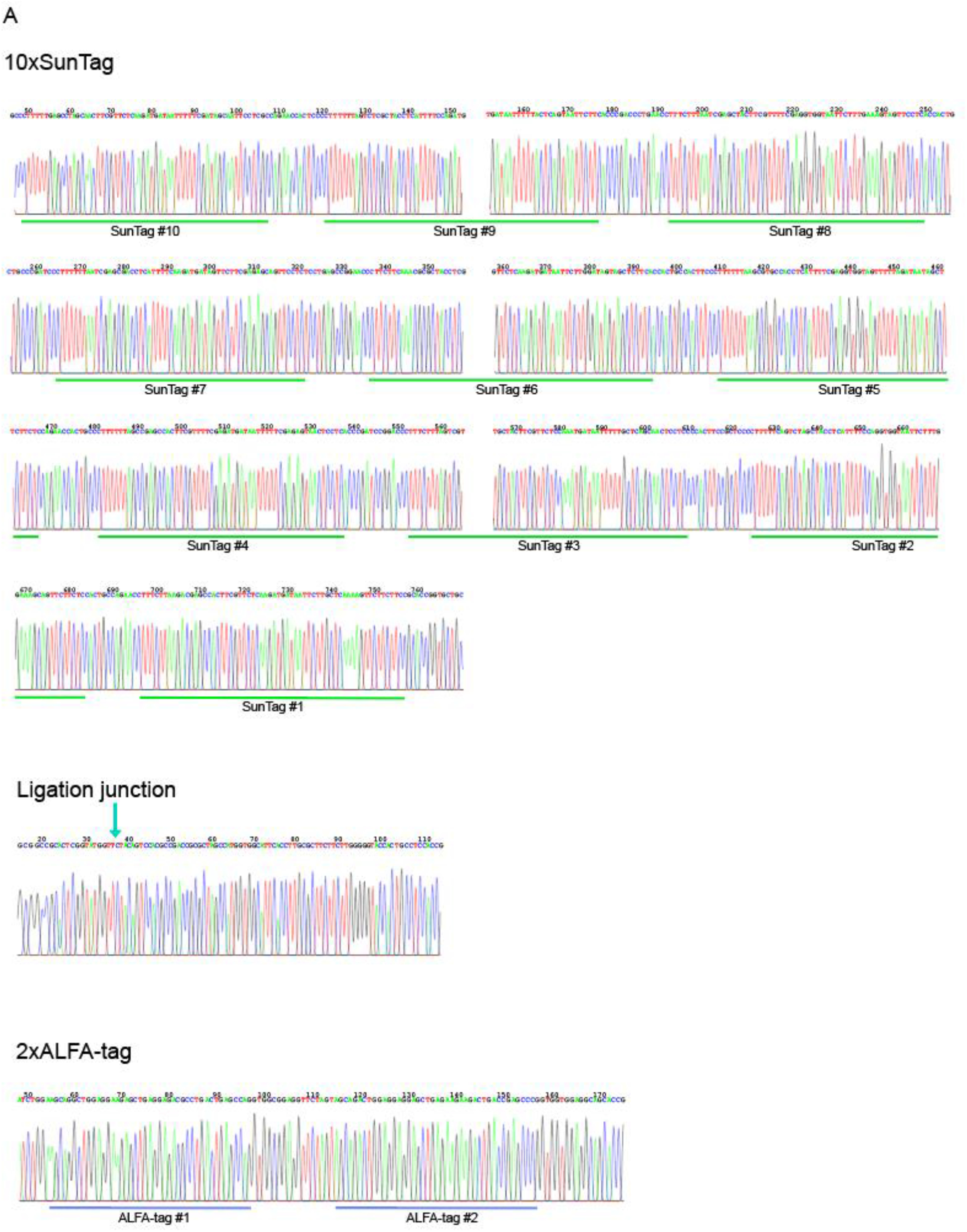
related to Figure 5. Validation of socRNA sequence. A) 10xSunTag socRNAs were sequenced using Sanger sequencing (See Methods). Three separate sequencing reactions were performed to sequence the 10xSunTag array (top), the circRNA ligation junction (middle) and the 2xALFA-tag sequence (bottom). Green lines underneath sequencing results indicate the position of the individual SunTag repeats, cyan arrow denotes the socRNA ligation site after circularization, and blue lines indicate the position of the ALFA-tag repeats. Note that only a single nucleotide was present in the sequencing reaction, indicating that the socRNAs expressed inc cells mostly have the same (correct) sequence.

**Figure S6.**
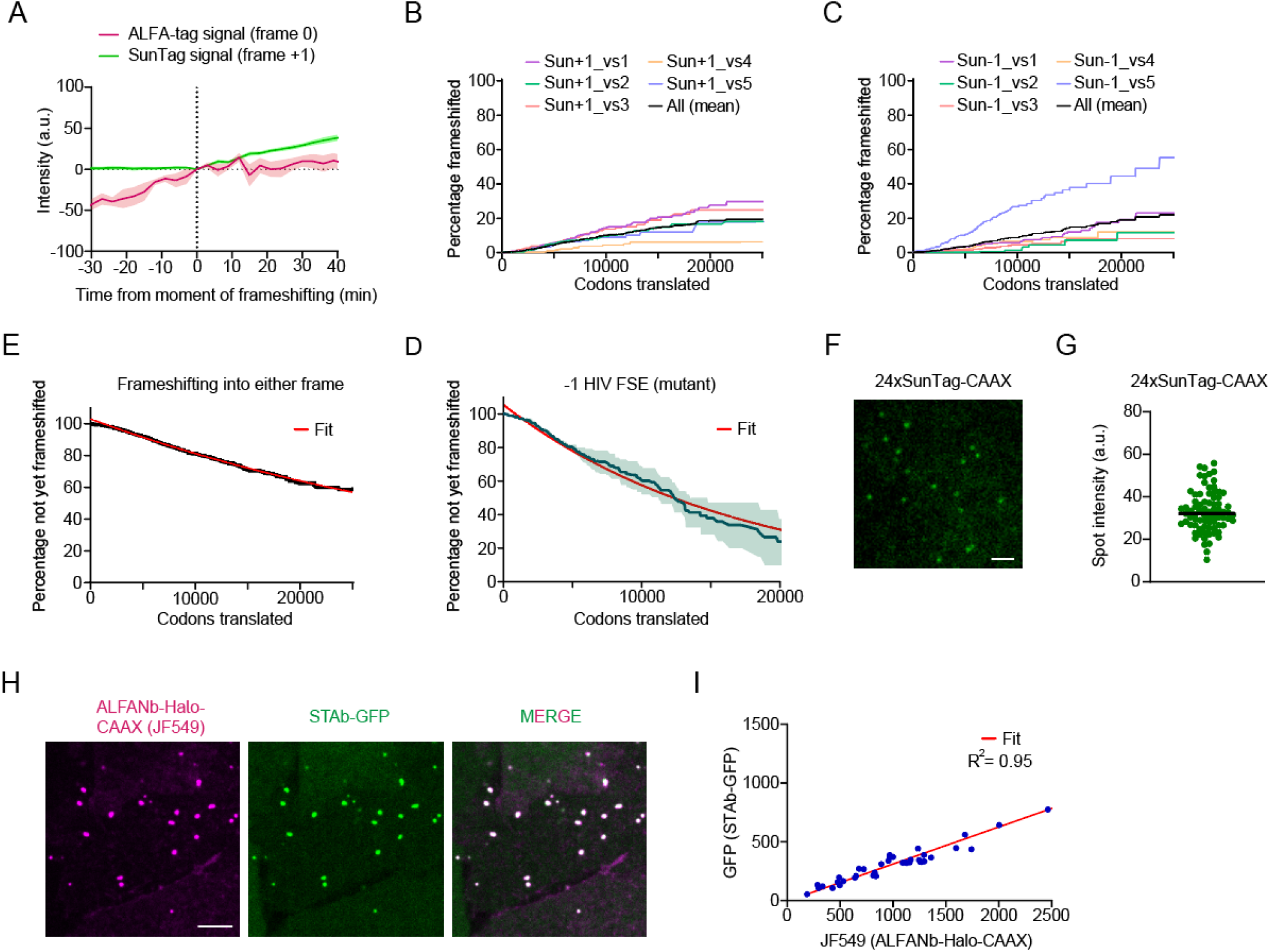
related to Figure 6. Controls for the socRNA frameshifting assay. A) Intensities from live-cell imaging of frameshifting into +1 frame were aligned to the moment of frameshifting. Lines indicate mean value and shaded regions indicate standard error of the mean. B-C) Kaplan-Meyer survival curve for socRNAs to measure frameshifting into the +1 frame (B) or the -1 frame (C) showing the total number of codons translated by ribosomes before frameshifting occurs (See Methods). D) Kaplan-Meyer survival curve (replotted from Figure 6G) showing frameshifting into either +1 or -1 frame. A single-exponential decay function (red line) was fitted to the data to calculate frameshifting rate per number of translated codons. E) Kaplan-Meyer survival curve showing the frameshifting rate into the -1 frame for a control reporter encoding a weak mutant of the HIV –1 programmed ribosomal frameshifting element. The red line indicates an exponential decay fit used to calculate frameshifting rate per number of translated codons. Lines indicate mean values and shaded regions indicate standard error of the mean. F-G) A construct encoding 24xSunTag-CAAX was expressed to determine the intensity produced by 24xSunTag epitopes. Line indicates median value. H-I) socRNA encoding 1xSunTag and 1xALFA-tag in the same reading frame was expressed to correlate the intensities of STAb-GFP and ALFANb-Halo. Scale bars, 2 μm (F), 5 μm (H). The number of experimental repeats and cells analyzed per experiment are listed in Table S1.

**Video S1 – related to Figure 1. Long-term imaging of single translating ribosomes using socRNAs**.

U2OS cells stably expressing STAb-GFP, ALFANb-CAAX, and tetR were transfected with a plasmid encoding a socRNA with 5xSunTag and 1xALFA-tag under the control of a doxoycline-inducible CMV promoter. SocRNA expression was induced by adding doxocycline to cells for 5 minutes, after which cells were washed and positions selected. Images were acquired every three min for four consecutive hours. Scale bar, 10 μm.

**Video S2 – related to Figure 1. Release of nascent chains by puromycin enables quantification of the number of ribosomes translating each socRNA**.

U2OS cells stably expressing STAb-GFP, ALFANb-CAAX, and tetR were transfected with a plasmid encoding a socRNA with 5xSunTag and 1xALFA-tag. Puromycin was added to socRNA-expressing cells at t=60 to induce nascent chain release and to allow scoring of the number of ribosomes per socRNA. Four separate movies from the same live-cell imaging experiment were combined into a single movie for side-by-side comparison of socRNAs translated by one, two, three or four ribosomes, respectively. Images were acquired every two minutes. Scale bar, 2 μm.

**Video S3 – related to Figure 6. Real-time imaging of frameshifting by single ribosomes**.

U2OS cells stably expressing STAb-GFP, ALFANb-Halo-CAAX, and tetR were transfected with a socRNA expression plasmid encoding 1xALFA-tag and 1xSunTag in two separate reading frames. Importantly, neither the ALFA-tag frame (frame 0) nor the SunTag frame (frame +1) contain a stop codon. Shown is a presentative movie of a ribosome undergoing frameshifting from frame 0 into the +1 frame. Puromycin was added at t=180 to release ribosome nascent chains. ALFA-tag and SunTag signal do not separate upon puromycin addition, indicative of a single, chimeric protein produced by ribosome frameshifting. Images were acquired every four minutes. Scale bar, 2 μm.

## Notes

### Competing Interest Statement

The authors have declared no competing interest.

